# Helicobacter pylori Senses Bleach as a Chemoattractant Using a Cytosolic Chemoreceptor

**DOI:** 10.1101/544239

**Authors:** Arden Perkins, Dan A. Tudorica, Manuel R. Amieva, S. James Remington, Karen Guillemin

## Abstract

The gastric pathogen *Helicobacter pylori* requires a non-canonical cytosolic chemoreceptor transducer-like protein D (TlpD) for efficient colonization of the mammalian stomach. Here we reconstituted a complete chemotransduction signaling complex *in vitro* with TlpD and the chemotaxis proteins CheW and CheA, enabling quantitative assays for potential chemotaxis ligands. We found that TlpD is selectively sensitive at micromolar concentrations to bleach (hypochlorous acid, HOCl), a potent antimicrobial produced by neutrophil myeloperoxidase during inflammation. Counterintuitively, HOCl acts as a chemoattractant by reversibly oxidizing a conserved cysteine within a 3His/1Cys Zn-binding motif in TlpD that inactivates the chemotransduction signaling complex. We found that *H. pylori* is resistant to killing by millimolar concentrations of HOCl and responds to bleach in the micromolar range by increasing its smooth swimming behavior, leading to chemoattraction to HOCl sources. We found that related protein domains from *Salmonella enterica* and *Escherichia coli* showed a similar reactivity toward bleach. We propose that this family of proteins enables host-associated bacteria to sense sites of tissue inflammation, a strategy that *H. pylori* uses to aid in colonizing and persisting in inflamed gastric tissue.

## INTRODUCTION

*Helicobacter pylori* is a bacterial pathogen and persistent colonizer of the human stomach and causes gastritis, ulcers, and stomach cancer (Amieva and Peek, 2016). The health burden caused by *H. pylori* is particularly large because it infects about half the world’s population, with nearly 100 % infection rates in some developing regions, and drug resistance to first line antibiotics is increasing (Amieva and Peek, 2016; Mégraud, 2012). Despite triggering a robust inflammation response and bursts of reactive oxygen species (ROS) from immune cells, *H. pylori* avoids clearance and persists for many decades (Ramarao et al., 2000). The chronic inflammation induced by *H. pylori* infection is thought to be a major factor in causing disease (Kozol et al., 1991). Not only is *H. pylori* not eradicated by inflammation, the pathogen in fact has been shown to navigate to sites of injury and is reported to use host iron extracted from blood hemoglobin and transferrin (Aihara et al., 2014; Husson et al., 1993; Tan et al., 2011; Worst et al., 1995).

*H. pylori* utilizes chemotaxis to seek sites optimal for growth and colonization within the hostile environment of the stomach (Johnson and Ottemann, 2018)(Fig. 1A). Bacterial chemotaxis is a well-studied phospho-relay system prevalent across bacteria and archaea, and functions through a conserved mechanism (Ortega et al., 2017). Chemoreceptor proteins typically possess a periplasmic ligand sensing domain to recognize small molecules and transduce signals across the inner membrane to a cytosolic coiled-coil region to regulate the autophosphorylation of the histidine kinase CheA with ATP (Hazelbauer et al., 2008)(Fig.1A). As the cellular pool of CheA-Pi is increased, the flagella rotors frequently reverse and alter the swimming bacterium’s trajectory, leading to chemorepulsion; decreases in CheA-Pi result in smooth swimming and chemoattraction (Fig. 1A). Chemoreceptors in *H. pylori* and other bacteria oligomerize to form trimers-of-receptor dimers (Fig. 1B) to build repeating hexagonal arrays that serve to amplify ligand-induced signals up to 50-fold, whereby a single activated receptor can initiate the signal transduction cascade (Briegel et al., 2012; Hazelbauer et al., 2008; Qin et al., 2017). The minimal core unit for signaling is thought to contain two trimers-of-receptor dimers, a CheA dimer, and 2-4 CheW scaffold proteins (Briegel et al., 2012; Cassidy et al., 2015; Haglin et al., 2017) (Fig.1B).

**Figure 1.**
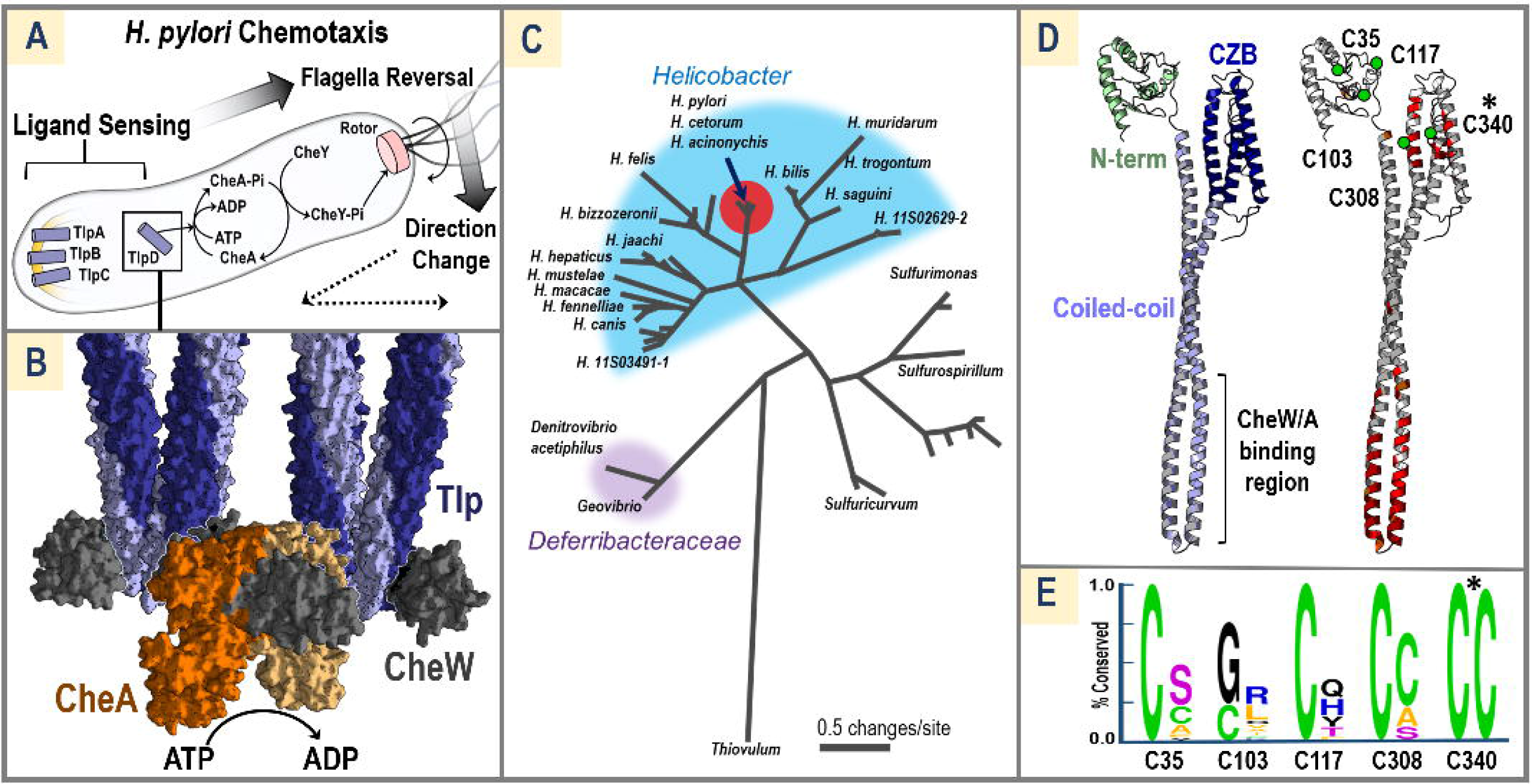
Conservation and Architecture of TlpD. (A) *H. pylori* inner membrane-bound (yellow arc) chemoreceptors TlpA-C, and cytosolic TlpD, control the autophosphorylation of CheA to CheA-Pi, which then transfers the phosphate to CheY, and CheY-Pi can interact directly with the flagella rotor (pink) to cause a temporary reversal in flagellar rotation. Flagella reversals cause direction changes in swimming trajectory (dotted lines). (B) Two sets of chemoreceptor trimers of homo dimers (light and dark blue, denoted as Tlp) associate with the scaffold protein CheW (grey) and a CheA dimer (light and dark orange) to form the core signaling unit and modulate CheA autophosphorylation (PDB code: 3ja6)(Cassidy et al., 2015). (C) A relatedness tree of TlpD protein sequences from *Helicobacter* (blue), *Deferribacteraceae* (purple), and other species. *H. pylori* TlpD (arrow) is nearly identical to species found in dolphins and cheetas (red dot). (D) A theoretical model of a *Hp*TlpD monomer constructed by i-Tasser with an N-terminal region of unknown structure (light green), the canonical coiled-coil domain of chemoreceptors that interfaces with CheW and CheA (light blue), and a chemoreceptor zinc-binding domain (CZB, dark blue). Sequence conservation amongst 459 TlpD homologues is mapped onto the model, with 100 % conservation highlighted in red, and >95 % conservation in orange. Cys residues present in the SS1 strain of *Hp*TlpD are noted with green circles. (E) Conservation of Cys residues in TlpD is shown with two Weblogo plots (Crooks, 2004) for TlpD Cys residues; the left shows conservation among all sequences of *Hp*TlpD (394 sequences, >92 % sequence coverage, >98 % sequence identity) and the right plot is conservation among non-*pylori* TlpD sequences (68 sequences, >91 % coverage, >40 % sequence identity). Only C340 is universally conserved among all TlpD homologues (noted with * in E and D).

*H. pylori* chemotaxis is mediated by four chemoreceptors, transducer-like proteins (Tlp) A-D (Fig. 1A). Environmental chemicals that elicit a chemoresponse by *H. pylori* include acid as a chemorepellent (Goers Sweeney et al., 2012; Huang et al., 2017), and urea as a chemoattractant (Huang et al., 2015), which both serve to direct *H. pylori* from the stomach lumen to the gastric epithelium. Experiments with Tlp genetic deletions have implicated chemoreceptors in cellular sensing processes, but only two chemoeffectors have been confirmed in biochemical experiments to be direct binding interactions between ligand and receptor: TlpB sensing of urea (Goers Sweeney et al., 2012; Huang et al., 2015) and TlpC sensing of lactate (Machuca et al., 2017), though the latter receptor is not conserved and apparently dispensable. Some progress has been made for understanding direct ligand sensing by TlpA, for which recent crystal structures suggest binding of a small hydrophobic ligand (Goers Sweeney et al., 2012). Sensing of pH remains more complicated to understand, as genetic knockouts indicate signals are integrated from TlpA, B, and D (Huang et al., 2017), and no direct sensing mechanism is known for TlpD.

Interestingly, it is the cytosolic chemoreceptor TlpD that is the most highly expressed in *H. pylori*, constituting about half of the total chemoreceptor pool (Fig. 1A)(Goers Sweeney et al., 2012). Such cytosolic or “soluble” chemoreceptors are widespread and common, accounting for approximately 15-percent of all bacterial and 45-percent of archaeal chemoreceptors, but their functions are mostly unknown and to date only a few cytosolic chemoreceptors have been mapped to their ligands (Collins et al., 2014). Like membrane-bound homologues, cytosolic chemoreceptors form nanoarrays (Briegel et al., 2014) and therefore represent an intriguing, fully soluble model system to better understand chemoreceptor function. No study has yet described the assembly of cytosolic signaling units in terms of the thermodynamic parameters or reaction kinetics for a cytosolic receptor with CheW and CheA.

A clear consensus on the molecular function of TlpD has remained elusive. Single receptor knockout strains of *H. pylori* show defects in colonization of the stomach antrum, a preferred niche for the bacterium and the region of highest inflammation, and of these, *tlpD* strains are by far the most impaired (Rolig et al., 2012). The first molecular study of TlpD revealed the protein to possess a novel chemoreceptor zinc-binding domain (CZB) that is prevalent among chemoreceptors and some diguanylate cyclases (Draper et al., 2011a). The domain uses a rare 3His/1Cys motif to coordinate the bound zinc with femtomolar affinity, and the zinc was unable to be removed even by the strong zinc chelator TPEN (Draper et al., 2011a). Shortly thereafter, crystal structures of a wild type and Cys→Ala mutant CZB domain were solved for DgcZ (previously called YdeH), an *E. coli* diguanylate cyclase, and those authors suggested CZBs function as zinc sensors, and showed for that protein that the zinc could be chelated by millimolar concentrations of EDTA (Zähringer et al., 2013). Additionally, studies using chemotaxis assays to monitor *H. pylori* swimming patterns have implicated TlpD as a redox sensor, suggesting it responds to extracellular sources of ROS. However, there are discrepancies in the literature regarding this point. Recent work reported TlpD-dependent chemorepulsion from paraquat-generated superoxide (Collins et al., 2016), but another study observed a response consistent with chemoattraction to superoxide (Behrens et al., 2016). Additionally, TlpD-dependent chemorepulsion from hydrogen peroxide (H_2_O_2_) was reported (Collins et al., 2016), but the data presented was at millimolar concentrations of H_2_O_2_ whereas *in vivo* concentrations are thought to rarely exceed the low micromolar range (Halliwell et al., 2000; Schröder and Eaton, 2008). TlpD sensing of ROS was also suggested to function as an “energy sensor” by responding to cytosolic oxidants produced through metabolism (Behrens et al., 2016; Schweinitzer et al., 2008). No signaling mechanism has yet been proposed or vetted that directly shows that TlpD is a redox or zinc sensor.

Here, we have taken advantage of the solubility of TlpD to reconstitute the full (TlpD, CheW, CheA) chemotaxis signaling complex *in vitro* to directly assay thermodynamic and kinetic parameters and test the effects of putative ligands on receptor signaling. We show TlpD is fully active and able to strongly promote CheA autophosphorylation even without the addition of any ligand. Our biochemical characterization of TlpD provides no evidence to support direct sensing of zinc, pH, hydrogen peroxide, or superoxide. Instead, we show that the strong oxidant hypochlorous acid (HOCl, bleach), the major oxidative product generated by neutrophilic inflammation, potently and reversibly inactivates the signaling complex through a universally conserved cysteine in the TlpD CZB to elicit a chemoattractant response, and that reactivity toward HOCl is a conserved feature of CZB domains. We demonstrate a mechanism by which the cysteine forms a redox “Cys-Zn switch” that is tuned to be highly reactive toward HOCl and much less reactive toward peroxide and superoxide. Lastly, we perform *in vivo* assays that show *H. pylori* tolerates millimolar concentrations of HOCl and uses TlpD for chemoattraction to bleach at physiologically-relevant concentrations. We propose this mechanism has evolved to facilitate *H. pylori* chemoattraction to sites of inflammation and persistence in neutrophil-rich gastric glands.

## RESULTS

### TlpD Homologues Possess a Universally Conserved Zn-binding Cysteine

To identify conserved regions important to TlpD function we performed a protein sequence BLAST search to retrieve putative homologues (Altschul et al., 1990). This search revealed that TlpD is almost exclusively found in the family *Helicobacteraceae* with most homologues being from mammal-associated strains, and a few TlpDs from reptile-associated strains (Fig. 1C). As expected, TlpD sequences from *H. pylori* were highly similar, with >98 % sequence identity across 394 isoforms in the non-redundant sequence database. To better understand sequence conservation on a structural level the i-Tasser program (Zhang, 2008) was used to generate a homology model of the receptor, as no experimental structure for TlpD is available (Fig. 1D). This model predicts three general domains: an N-terminal region of low sequence conservation and unknown function, the canonical chemoreceptor coiled-coil that interacts with CheW and CheA, and a C-terminal chemoreceptor Zn-binding domain (CZB). Mapping positions of high conservation onto the model reveals that the CheW/CheA interface (common to all chemoreceptors) is highly conserved, as are positions in the CZB domain near the Zn-binding residues (Fig. 1D). *H. pylori* TlpD is unusual for a chemoreceptor, as it contains an abundance of Cys residues (4-5 depending on the strain): C35, C103, C117, C308, and C340 (Fig. 1D). Four of the Cys positions are conserved among *H. pylori* TlpDs, but only C340 within the CZB domain is conserved across all homologues (Fig. 1E).

### Reconstituted Chemoreceptor Signaling Complex Retains Function In Vitro

Recombinant *H. pylori* TlpD, the scaffold protein CheW, and the histidine kinase CheA were expressed and purified for use in functional assays to directly test the effects of mutations and potential ligands using radio-ATP labeling to monitor autophosphorylation of CheA (see Method Details). In this assay, activation of CheA increases CheA autophosphorylation which promotes swimming reversals and chemorepulsion; decreasing CheA activity results in smooth-swimming and chemoattraction (Fig. 1A). Autophosphorylation kinetics of *Hp*CheA alone revealed a K_M_ for ATP of 136 μM, similar to the ~300 μM K_M_ reported for *E. coli* CheA, however the basal level of K_cat_ was exceptionally slow at 8.25 × 10^−3^ min^−1^ (compared to 1.56 min^−1^ for *E. coli*)(Tawa and Stewart, 1994)(Fig. 2A). We next determined the concentration of TlpD required to achieve full dimerization, since the receptor dimer is the core building block for the chemotaxis signaling complex (Haglin et al., 2017). The TlpD dimer K_D_ was found by analytical ultracentrifugation to be 150 nM, consistent with fluorescence anisotropy experiments that showed at approximately 16 μM and above, TlpD is fully dimerized (Fig. 2B). Based on these results, assays with TlpD were generally run at concentrations of 20 μM or higher so that the properties of the physiologically relevant dimer were observed rather than the inactive monomer. Addition of stoichiometric concentrations of TlpD, CheW, and CheA resulted in synergistic activation of the chemotaxis complex; CheW was able to partially activate CheA alone, and activation was further increased with the addition of TlpD (Fig. 2C). These results confirmed that the recombinant system was functional and activated CheA in a manner similar to other described chemotaxis systems (Gegner et al., 1992).

**Figure 2.**
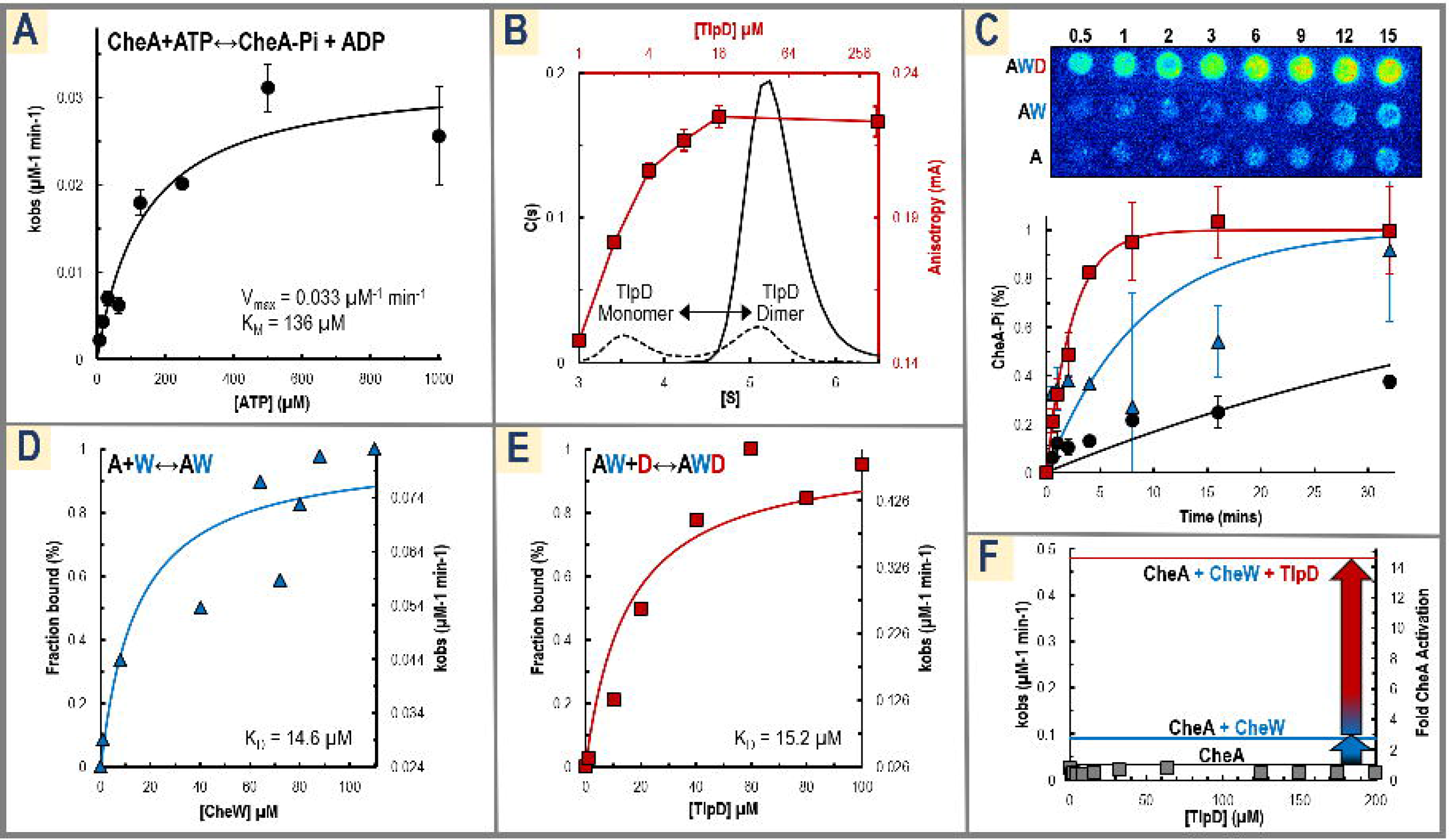
TlpD and CheW Synergistically Activate CheA. (A) Kinetics of *Hp*CheA with varying concentrations of ATP are shown. Experiments were conducted in triplicate with 4 μM CheA and varying amounts of ATP, in 50 mM Tris pH 7.5, 100 mM NaCl, 10 mM MgCl_2_. The average k_obs_ for three replicate time courses are shown at various [ATP] (black dots), error bars are the sample standard deviation, and these measurements are fit to the Michaelis-Menton curve (black line). (B) Representative analytical ultracentrifugation data (black axes) are shown for TlpD at 1 μM (black dotted black line) and 4 μM (black solid line) in PBS buffer pH 7 with 1 mM TCEP. Peaks corresponding to the TlpD monomer and dimer occur near 3.5 [S] and 5.2 [S], respectively, with a K_D_ of 150 nM. Shown in red on secondary axes are fluorescence anisotropy data for a titration of TlpD under identical conditions with experiments run in triplicate. (C) Shown top are representative raw data from radio-ATP labeling experiments of 15-minute reactions with CheA alone (“A”), and additions of CheW (“AW”) and TlpD (“AWD”). Below are reactions of 1 mM ATP and 4 μM CheA (black circles), +8 μM CheW (blue triangles), and +8 μM CheW, 24 μM TlpD (red squares) run in triplicate and fit to a pseudo-first order reaction curve (solid lines). (D) CheW was titrated against 4 μM CheA and resulting k_obs_ measurements were fit to a binding isotherm to obtain a kinetically-defined K_D_ of 14.6 μM for the CheA↔CheW interaction. (E) TlpD was titrated against 4 μM CheA and 40 μM CheW and fit to a binding isotherm as in C to approximate the thermodynamics of the AW↔D interaction to have a K_D_ of 15.2 μM. (F) A titration of TlpD against 4 μM CheA shows no activation (gray squares). For CheA in the presence of saturating [CheW] a 2.7-fold activation occurs (blue line), and with saturating [CheW] and [TlpD] this is increased to a 14.6-fold activation (red line) over CheA alone (black line). See Table S1 for a summary of reaction parameters.

We next leveraged this system to obtain kinetically-defined thermodynamic binding constants for the components and determine the required order of complex assembly. Titration of CheW against CheA revealed a K_D_ of 14.6 μM for formation of the CheW-CheA complex, similar to that of *E. coli* at 17 μM (Gegner and Dahlquist, 1991), and maximal activation of 2.7-fold (Fig. 2 D). Titration of TlpD against CheW-CheA showed a K_D_ of 15.2 μM, and maximal 14.6-fold activation of the complex (Fig. 2E). Previous work has shown for *E. coli* that CheW is required for the receptor to activate CheA (Boukhvalova et al., 2002), but a recent study suggested TlpD could activate CheA independently (Abedrabbo et al., 2017). However, that study used non-stoichiometric ratios for the chemotaxis components and only 2 μM TlpD, and based on our measured K_D_ of 150 nM approximately 20 % of the receptor would be expected to be in its inactive monomer form. Direct activation of CheA by TlpD was tested with a titration of TlpD against CheA without CheW, but in our hands no activation occurred even at 200 μM receptor (Fig. 2F). Instead, the data supports a sequential activation of the complex whereby CheA and CheW first associate to achieve 18-percent activation, and then TlpD can bind and promote full, 100-percent activation of the chemotaxis complex (Fig. 2F). To our knowledge this is the first characterization of the thermodynamic and kinetic parameters of a cytosolic chemotaxis complex and indicates in this case that the chemoreceptor TlpD is “on” by default, and able to activate CheA even without the addition of other ligands. Kinetic and thermodynamic parameters are summarized in Table S1.

### TlpD is Not Directly Sensitive to Exogenous Zn^++^, pH, Hydrogen Peroxide, or Superoxide

Two lines of evidence led us to hypothesize the CZB domain might in fact use zinc as a cofactor in ligand sensing, rather than directly sense Zn^++^. First, TlpD binds zinc with femtomolar affinity (Draper et al., 2011b), which seems inconsistent with a sensor that can be activated and deactivated by exogenous zinc to regulate bacterial swimming, as other bacterial chemoreceptors typically exhibit ligand affinity in the micromolar range (Clarke and Koshland, 1979; Goers Sweeney et al., 2012; Ortega et al., 2017). We examined whether it was possible to remove zinc from TlpD by chelating agents to test if a reversible zinc-sensing function were feasible. The fluorescent probe Zinpyr-1, which binds zinc with nanomolar affinity, was incubated with TlpD, but no chelation was observed (Fig. 3A-B). We also heat denatured TlpD, and although a small amount of zinc was released, most was retained by the protein, suggesting that even when unfolded the zinc is sufficiently buried to prevent removal (Fig. 3A). Previous ICP mass spec analysis of TlpD showed the protein is fully zinc-loaded following purification (Draper et al., 2011a), but to verify our purified protein followed suit, Zn was added at 0.25x, 0.5x, and 1x (24 uM) zinc to the functional assay. The addition had no effect on autophosphorylation rates except at higher concentrations (far beyond physiologically-relevant), where proteins were observed to precipitate and be inactivated (Fig. 3C). We also explored a potential role of the zinc binding core in direct pH sensing. If TlpD regulation of CheA is sufficient to stimulate chemorepulsion from acid and chemoattraction to basic pH, this would be reflected in our activity assay as decreasing CheA autophosphorylation as a function of increasing pH, however functional assays across different pHs did not replicate *in vivo* responses (see Method Details and Fig. S1). Activity was similar across pH 6.6-7.8, the normal range that TlpD would be expected to experience within the well-buffered cytosol (Stingl et al., 2001). Together, these data led us to conclude that although TlpD is important for acid sensing *in vivo* the receptor does not directly sense pH, and in terms of zinc binding, the protein as purified is fully zinc-loaded, binds zinc with extremely high affinity and does not release it, and is therefore unlikely to directly detect exogenous zinc as a chemotaxis ligand.

**Figure 3.**
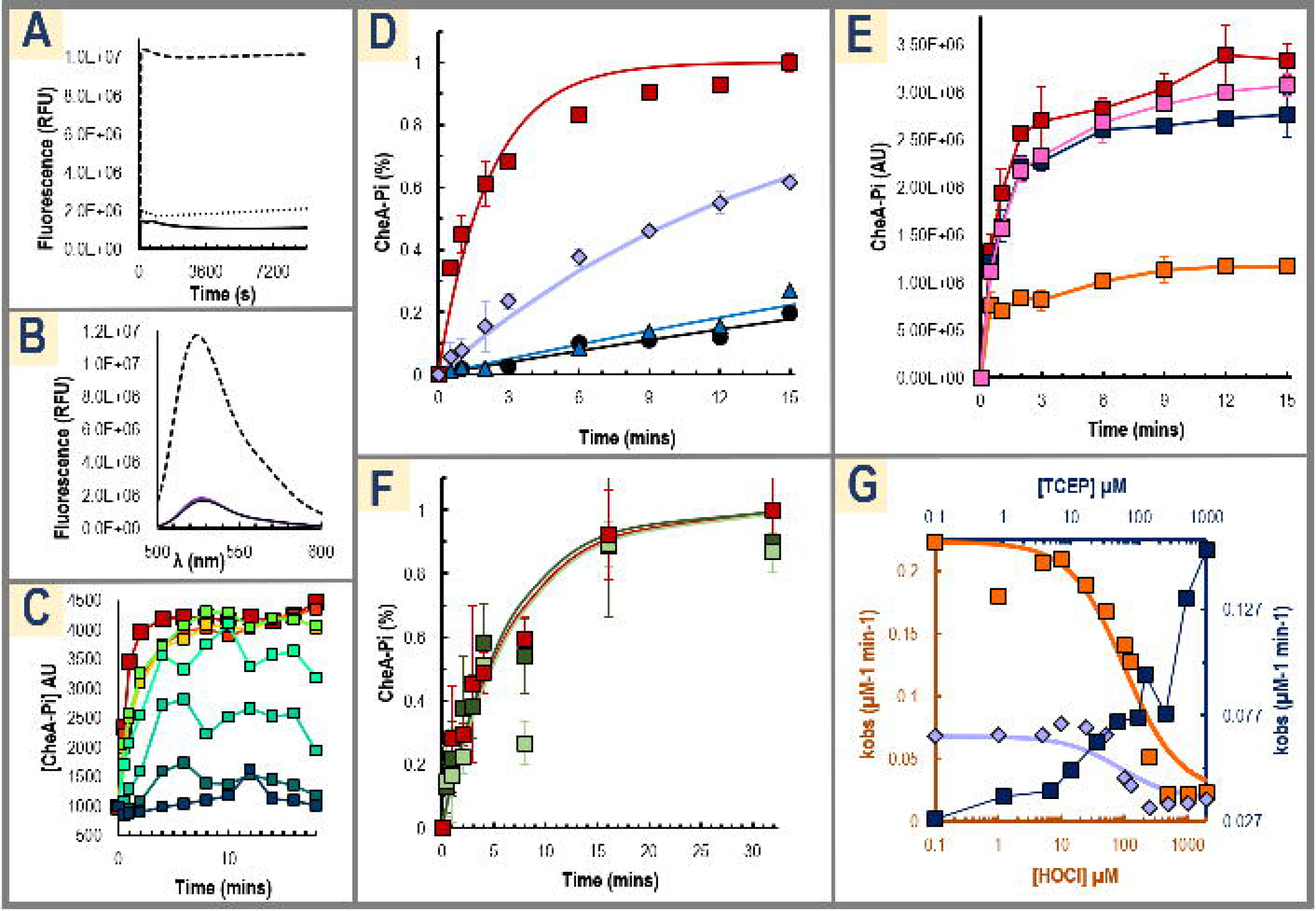
Dissecting TlpD Activation of CheA. (A-C) Three experiments are shown that demonstrate Zn binds tightly to TlpD and cannot be extracted. (A) Time courses monitoring fluorescence (ex. 429/em. 527 nm) for 50 μM Zinpyr-1 with either 50 μM zinc acetate (black dashes), 50 μM TlpD (black), or 50 μM heat-denatured TlpD (black dotted) shows no significant chelation. (B) Shown is a fluorescence spectrum (ex. 492 nm) using 50 μM of the Zn-chelating probe Zinpyr-1. A positive control adding 300 μM zinc acetate shows a strong increase in fluorescence at 547 nm (black dashes), but addition of 30 μM TlpD for 10 mins (purple) does not increase fluorescence over Zinpyr-1 alone (black, closely overlays with purple). (C) A series of functional assays with addition of 0 (red), 0.25x (orange), 0.5x (yellow), 1x (lime green), 2x (teal), 4x (cyan), 8x (dark green), and 16x (dark blue) zinc sulfate relative to [TlpD]. (D) Reaction time courses of reconstituted signaling complex are shown using either wild type TlpD (red squares) or C340A (violet diamonds), with single time courses for CheA (black circles), and CheA, CheW (blue triangles) for reference. (E) Hypochlorous acid (HOCl), but not H_2_O_2_ or reduction, directly alters CheA activation. Shown are functional assay time courses following 1-H pre-treatments with either buffer (red), 500 μM H_2_O_2_ (pink), TCEP (dark blue), or buffered sodium hypochlorite (NaOCl, orange)(n=3). (F) Paraquat does not directly alter activation of CheA by TlpD. Functional assays are shown with either 1-H pretreatment with buffer (red), 10 μM paraquat (light green), or 100 μM paraquat (dark green). (G) A series of functional assays titrating with NaOCl is shown with k_obs_ values from reaction time courses plotted against [NaOCl] on a log scale (orange boxes). Fitting these data to a binding isotherm yields a K_1/2_ for inhibition of TlpD-activation of CheA activity by NaOCl of 100.5 μM (orange line). An identical titration was performed for the C340A mutant (violet diamonds and line). Recovery of inactivated complex with wild type TlpD by reduction with TCEP is shown in dark blue, plotted on a secondary axis. Complex was pre-treated for 1-H with 125 μM NaOCl, and then treated with varying [TCEP] for 30 minutes. See also Fig. S1 for analysis of TlpD activity at various pH.

The second line of evidence suggesting zinc as a cofactor comes from the crystal structure of the *E. coli* CZB in which the cysteine residue equivalent to the conserved TlpD C340 was mutated to an alanine (PDB code: 4h54). In this structure, the zinc remained bound by the 3-His core even without coordination by the Cys (Zähringer et al., 2013), hinting the Cys could reversibly bind and detach from the zinc in response to some molecular cue. To test if C340 is important for promoting CheA autophosphorylation, activity in a functional assay using either wildtype TlpD or a C340A mutant was compared, and the mutation was found to cause a 6.8-fold loss in CheA activation, supporting that the Cys is required for full activation by TlpD (Fig. 3D). To test if C340 might directly sense cellular oxidants, functional assays were performed with 500 μM H_2_O_2_ or 500 μM of the reductant TCEP but no change in activity was observed (Fig. 3E), and addition of paraquat at concentrations used in previous *H. pylori* chemotaxis assays (Behrens et al., 2016) also did not alter activation of the complex, suggesting TlpD does not directly sense superoxide (Fig. 3F).

### TlpD Directly Senses Hypochlorous Acid Through Oxidation of a Conserved Cysteine

Though previous work had tested TlpD-dependent sensing of hydrogen peroxide and superoxide (Behrens et al., 2016; Collins et al., 2016), and chemoattraction to sites of gastric injury has been shown *in vivo* (Aihara et al., 2014), no study had yet analyzed whether TlpD might respond to the inflammation oxidant HOCl. Addition of 500 μM HOCl to reconstituted complex dramatically decreases CheA autophosphorylation to the rates observed for CheW and CheA alone (Fig. 3E), and a titration with HOCl revealed a K_1/2_ of approximately 100 μM (Fig. 3G). A similar titration using the C340A mutant exhibited largely diminished activity even without the addition of HOCl, indicating the mutation may effectively mimic the “off” state of the receptor (Fig. 3G). In these assays no protein precipitation was observed, but because HOCl is known to oxidatively damage proteins and cause aggregation (Mahawar et al., 2011), HOCl-oxidized samples were tested for resurrection by the reductant TCEP, which was shown to effectively reactivate the complex (Fig. 3G).

These data suggest the mechanism of HOCl sensing involves oxidation of TlpD C340 to a cysteine sulfenate, an oxidation product commonly formed in redox systems that utilize a redox-active cysteine, and which is readily reversible by small molecule and protein reductants such as the thioredoxin/thioredoxin reductase system present in *H. pylori* (Poole and Nelson, 2008; Windle et al., 2000). Close analysis of the *E. coli* CZB structure revealed a small channel that allows solvent to access the cysteine, which has not been previously noted (Fig. 4A-B). The C340 S_γ_ is partially solvent exposed at the base of this cavity, reminiscent of an enzyme active site suitable for binding a small ligand. There, two water molecules mimic the approximate positions for an oxygen and chloride of a hypochlorite molecule and are coordinated by the backbone NH of the following helical turn to be well-positioned to react with the C340-S_◻_ (Fig. 4B-C). To determine if C340 can react with HOCl and become oxidized to a cysteine sulfenate, we treated wild type TlpD and C340A protein with HOCl and monitored formation of cysteine sulfenate by dimedone adduction and western blotting (Seo and Carroll, 2009). The results show potent formation of cysteine sulfenate for the wild type protein, but almost none for the C340A mutant (Fig. 4D). This indicates not only that HOCl can access and directly oxidize C340 in solution, but also that C340 is tuned to be highly reactive with HOCl, as it is approximately 50-fold more readily oxidized than the four other TlpD Cys residues combined (Fig. 4D). Titration data exhibit a sigmoidal response and are fit well with a Hill Coefficient of n=2, suggesting positive cooperativity occurs across the receptor homodimer (Fig. 4D). As a secondary test of direct oxidation of C340 by HOCl we performed mass spectrometry of reduced, H_2_O_2_-treated, and HOCl-treated TlpD to look for the formation of oxygen adducts. The spectra indicated little oxidation of C340 occurred for the reduced and peroxide-treated samples, but spectra of HOCl-treated TlpD showed C340 to be completely oxidized to sulfinate and sulfonate forms (Fig. 4E and Fig. S2A-C). Similar “over-oxidation” in the absence of reductant in *in vitro* assays is seen to occur for other proteins with strongly redox-active Cys residues (Perkins et al., 2013, 2015), but bacteria generally lack the capacity to reduce sulfinate and sulfonate forms (Perkins et al., 2014), so we expect that cysteine sulfenate is the oxidation state of physiological relevance. Lastly, we tested two other representative CZBs, the CZB domain of the *E. coli* diguanylate cyclase DgcZ and the *Salmonella enterica* chemoreceptor McpA. We found them to be similarly reactive toward HOCl as TlpD, suggesting HOCl-sensing is a conserved function of CZB domains (Fig. S2D).

**Figure 4.**
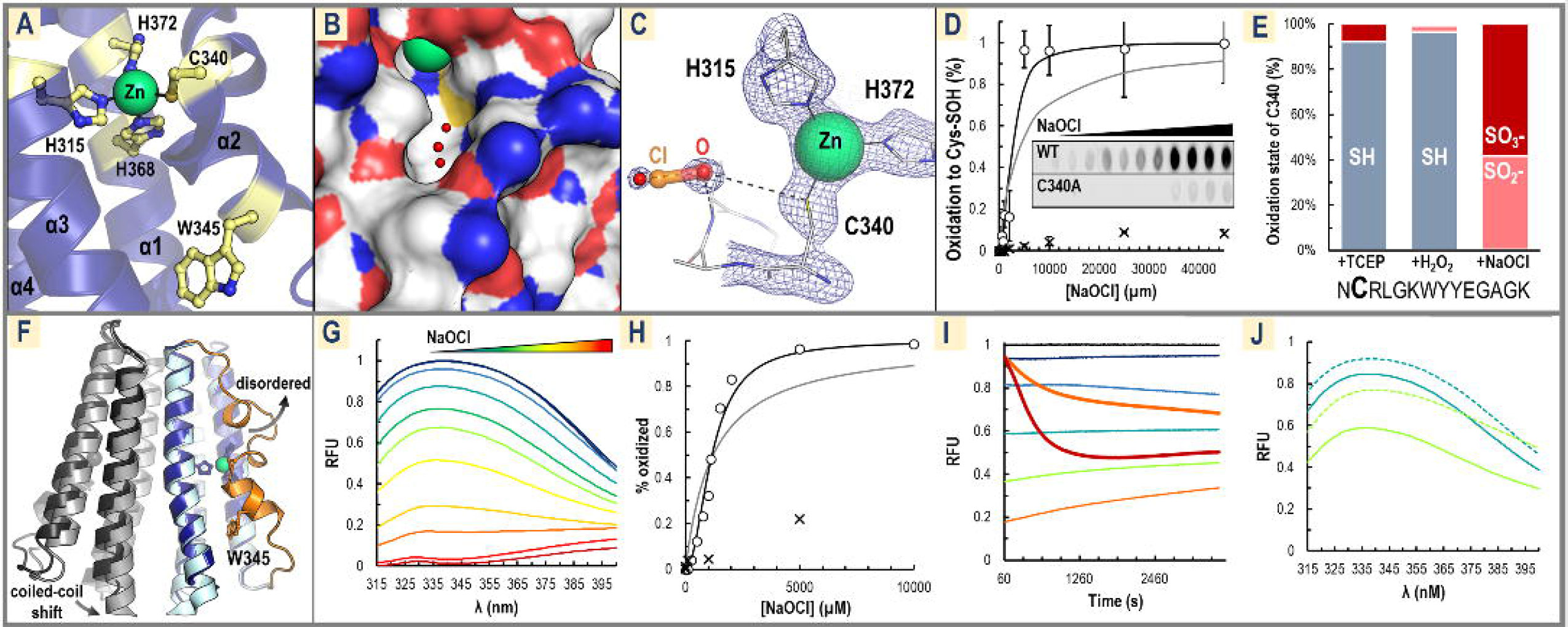
TlpD Senses HOCl by Reversible Oxidation of C340. (A) The crystal structure of the *E. coli* CZB domain is shown, with the conserved residues of the Zn-binding core of CZB and nearby W345 highlighted in gold (PDB code: 3t9o, numbering is for corresponding residues in *Hp*TlpD based on sequence alignment). (B) The molecular surface in the same view as in A (sulfur is yellow, nitrogen blue, oxygen red, carbons white, zinc green) showing C340 is accessible to solvent. Three waters in the experimental structure are shown as small red spheres. (C) A close-up view of the CZB crystal structure is shown where two water molecules bind in the pocket near C340, mimicking the predicted position of hypochlorous acid (O and Cl labeled and shown as transparent sticks). 2fo-fc electron density is shown as blue mesh, contoured at 1.5 σ. (D) Oxidation of 20 μM wildtype TlpD (open circles) or C340A mutant (crosses) to form cysteine sulfenate (Cys-SO^−^) is shown at various HOCl treatments and monitored using an anti-cysteine sulfenate antibody. Experiments were run in triplicate and representative raw data is shown inset, with the image length reduced by 50 %. Fits of the wild type data to the Hill equation using either a coefficient of 1 (gray line) or 2 (black line) are shown. See Fig. S2D for similar experiments with the CZB domain of *E. coli* DgcZ and *Salmonella* McpA. (E) Data from MS/MS experiments show 1 mM HOCl potently oxidizes C340, but not 1 mM H_2_O_2_. The 14-mer peptide containing C340 (noted in bold) was identified and found to be modified either by cysteine alkylation (grey, SH, indicating no oxidation), oxidation to a cysteine sulfinate (peach, SO ^−^), or oxidation to a cysteine sulfonate (dark red, SO ^−^). See also Fig. S2A-C and Method Details. (F) Shown is an overlay of the wild type *E. coli* CZB homodimer (dark blue, PDB code: 3t9o) and Cys→Ala mutant (light blue, PDB code: 4h54) crystal structures. Without the Cys-Zn bond, the region containing C340 and W345 (orange) locally unfolds and becomes disordered and solvent exposed. This structural rearrangement also promotes a ~3 Å shift in its dimer partner (wild type dark gray, Cys→Ala light gray) in the helix adjoining the coiled-coil that may propagate to the CheW/A interface to deactivate the signaling complex. (G) Intrinsic protein fluorescence excited at 295 nm for a titration of NaOCl against 20 μM TlpD is shown. Treatments were for 30 minutes with 20 μM TlpD in PBS pH 7 with addition of 0 (black), 100 μM (dark blue), 250 μM (light blue), 500 μM (teal), 750 μM (green), 1 mM (lime green), 1.1 mM (yellow), 1.5 mM (light orange), 2 mM (orange), 5 mM (red), or 10 mM (dark red) NaOCl. (H) Data from F is quantified with the RFU at 340 nm at each [NaOCl] fit to the Hill equation with a coefficient of 1 (gray) or 2 (black). Representative equivalent experiments performed with C340A are shown as crosses. (I) Time courses comparing fluorescence quenching at 340 nm (excited at 295 nm) of 20 μM TlpD by peroxide (two thick lines) and NaOCl (five thin lines) at concentrations of 0, 100 μM, 250 μM, 500 μM, 1 mM, 2 mM, and 5 mM, colored as in C. (J) TlpD fluorescence quenched by HOCl-treatment can be recovered by reduction. 20 μM TlpD samples were pre-oxidized (solid lines) with HOCl at concentrations of 500 μM (blue) and 1 mM (lime) for 30 minutes prior to reduction by equimolar concentrations of TCEP for 30 minutes (dashed lines).

A comparison of the wild type and Cys→Ala CZB crystal structures shows that without the Cys-Zn bond, the CZB undergoes a large structural shift involving 22 residues whereby the helix containing the cysteine locally unfolds and becomes disordered (Fig. 4F). We speculated that oxidation of C340 could similarly cause C340 to detach from the zinc site and promote a conformational change to transduce the signal by shifting the upstream coiled-coil region (Fig. 4F). Fortuitously, TlpD contains a single conserved Trp residue (W345), located near C340, that participates in this conformation change (Fig. 4A,F). Therefore, fluorescence of W345 was used as a native probe to detect the receptor’s conformation change in response to C340 oxidation. Strong Trp fluorescence is observed in untreated TlpD samples and is effectively quenched by HOCl concentrations in the μM-mM range, supporting that oxidation induces a conformation change like that observed in the crystal structures (Fig. 4H). Similar to the apparent cooperativity observed for oxidation of C340 (Fig. 4D), the conformation change exhibits a sigmoidal response (Fig. 4H). The C340A mutant is about 5-fold less sensitive to HOCl treatment than wild type, showing this conformation change is dependent on the presence of C340 (Fig. 4H). To ensure the loss in fluorescence was not due to protein denaturing, a similar titration was monitored by circular dichroism, and shows TlpD remains folded up to at least 10 mM HOCl (Fig. S2E). Trp fluorescence quenching time courses revealed HOCl reacts to completion within the sample mixing time (< 60 s), whereas quenching by hydrogen peroxide requires 6 to 7-fold higher concentrations and is very slow, with the reaction requiring approximately 20 minutes to reach completion (Fig. 4I). Quenched fluorescence of oxidized TlpD was able to be recovered by reduction with TCEP (Fig. 4J).

### Hypochlorous Acid Decreases Helicobacter pylori Swimming Reversals

HOCl can reach concentrations as high as 5 mM at sites of inflammation (Weiss, 1989), and so prior to investigating *H. pylori* sensing of HOCl *in vivo* we first determined the range of HOCl concentrations *H. pylori* can survive and remain motile. We used a video tracking assay to measure swimming velocities in response to acute treatments of bleach in the absence of chemotaxis for an *H. pylori cheA* mutant (strain G27), which is motile but cannot respond to chemotaxis signals, and a motile *E. coli* lacking all chemoreceptors (strain UU1250) (Fig. 5A). We found that *H. pylori* is extremely resilient against HOCl treatment, with full loss of motility observed only for treatments exceeding approximately 7 mM, whereas *E. coli* lost motility in treatments exceeding 2 mM bleach (Fig. 5A).

**Figure 5.**
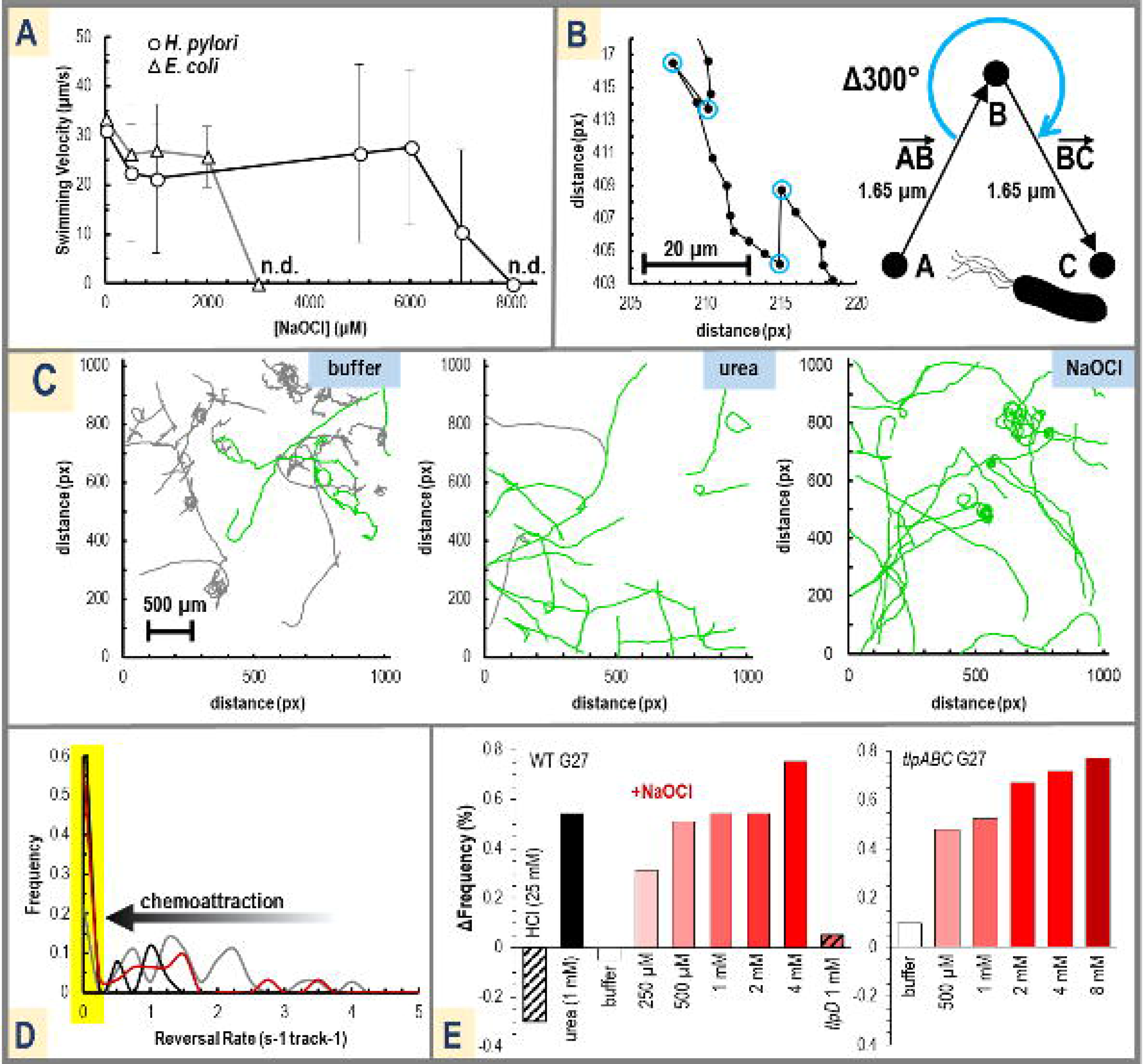
*H. pylori* Swimming Reversals are Decreased by Bleach Treatment. (A) *H. pylori* motility is more robust against bleach than *E. coli*. Shown is the mean swimming velocity versus bleach treatment using *E. coli* strain UU1250 (triangles, gray line) and *H. pylori cheA* (circles, black lines). The concentration of bleach at which no motile bacteria were detected is noted with “n.d.” Error bars shown are the standard deviation from the mean. Experiments were performed in triplicate, with n-values of >30 bacteria tracks (see Method Details). (B) Shown left is a representative swimming track for *H. pylori* with each dot corresponding to the bacteria’s position every 40 ms. Reversals are highlighted as blue circles. Shown right is the definition of a reversal we apply for quantification. (C) Representative swimming tracks of wild type *H. pylori* are shown following the addition of tris buffer, 1 mM urea, or 2 mM buffered NaOCl. Smooth-swimming tracks with reversal rates < 0.25/s are highlighted in green; others are gray. Note that these figures are intended to visualize the swimming behavior of the bacterial swarm but that individual reversals cannot be seen at this scale. (D) A histogram of reversal rates is shown for the data in C, with individual tracks binned by average reversal rate. Chemoattraction is visualized as a shift in the population to lower reversal rates. The untreated population (see Fig. S3F) and buffer-treated swarms (grey) exhibit approximately 20 % smooth swimming. The addition of 1 mM urea (black) or 2 mM NaOCl (red) shifts the population to be approximately 60 % smooth swimming (<0.25 reversals/s, highlighted in yellow). (E) Shown left are the changes in smooth swimming population (yellow box in D) of wild type swarms treated with the chemorepellent HCl (25 mM, black stripes, the chemoattractant urea (1 mM black), buffer (white), and various concentrations of NaOCl (red), and a *tlpD* mutant cotreated with 50 mM HCl and 1 mM NaOCl (red and black stripes). Shown right are treatments with a triple knockout *tlpABC* mutant. See Fig. S3 and Method Details for additional information.

We tested if *H. pylori* senses HOCl as a chemoattractant by observing bacteria swimming behavior following HOCl exposure. In this assay chemorepellants such as acid activate chemoreceptors and stimulate a buildup in CheA-Pi that results in more frequent flagella reversals and direction changes that here we refer to as “reversals” (Fig. 5B); conversely, chemoattractants inactivate chemoreceptors and reduce cellular CheA-Pi to promote smooth swimming (Collins et al., 2016). Analysis of wildtype strain G27 *H. pylori* in growth media showed a diverse range of reversal rates, with about 20 % of the tracks exhibiting smooth swimming (green tracks in Fig. 5C, Fig. S3). Urea was used as a positive control for chemoattraction (Huang et al., 2015), with addition of 1 mM urea shifting the swimming behavior to be approximately 75 % smooth swimming (Fig. 5C, Fig. S3B). The addition of 2 mM HOCl similarly decreased reversals and increased smooth swimming to approximately 70 %, consistent with chemoattraction behavior (Fig. 5C, Fig. S3).

Shifts to smooth swimming in terms of reversal rate histograms can be visualized as an increase in the 0-0.25 reversal/second bin (Fig. 5D). Using this metric, the frequency of smooth swimming was compared across controls and several concentrations of HOCl treatments. Treatments with bleach in the range of 250 μM to 4 mM resulted in a dose-dependent shift to smooth swimming behavior (Fig. 5E). Similar assays were performed with a *tlpD* mutant in a background of acid to promote enough reversals for detection, and no shift in swimming behavior was seen for bleach treatments (Fig. 5E, Fig. S3G and Method Details). HOCl treatments of a *tlpABC* mutant, which possesses TlpD as its sole chemoreceptor, resulted in a shift to smooth swimming similar to wild type (Fig. 5E, Fig. S3D-E). For both wild type and *tlpABC* 500 μM bleach was sufficient to elicit a smooth swimming phenotype equal to that of 1 mM urea (approximately 50 % increase in the smooth swimming frequency), and the entire population could be converted to smooth swimming with 4 mM HOCl, consistent with a chemoattraction response (Fig. 5E).

### Helicobacter pylori is Attracted to Hypochlorous Acid

Although the swimming assay provides useful insight into how bacterial swimming is altered by a chemoeffector, it is a proxy for the cellular levels of CheA-Pi and does not directly assay chemoattraction in terms of localization of bacteria toward a point-source. HOCl degrades at room temperature and is reactive with a variety of organic molecules, so commonly used point-source chemotaxis experiments such as capillaries and soft agar plates were not feasible. Instead, we utilized an assay in which a highly-calibrated micropipette can continuously deliver fresh treatment solution to establish a microgradient (Howitt et al., 2011; Huang et al., 2015). We tested wild type *H. pylori* strains PMSS1 and G27 for bleach chemoattraction and indeed found a rapid increase in the bacteria localization to a point-source of 10 mM HOCl, with the bacteria visible in frame after 30 s of treatment increasing by 100 and 75 %, respectively (Fig. 6A, Fig. S4A-B). HOCl chemoattraction was eliminated for a *tlpD* G27 mutant, indicating that TlpD is required for HOCl sensing (Fig. 6C-D, Fig. S4B). A weaker chemoattraction was retained for a *tlpABC* G27 mutant with an increase in bacteria of approximately 25 % over buffer after 30 s, suggesting TlpD is required and sufficient for bleach chemoattraction (Fig. 6 C,E, Fig. S4B-C).

**Figure 6.**
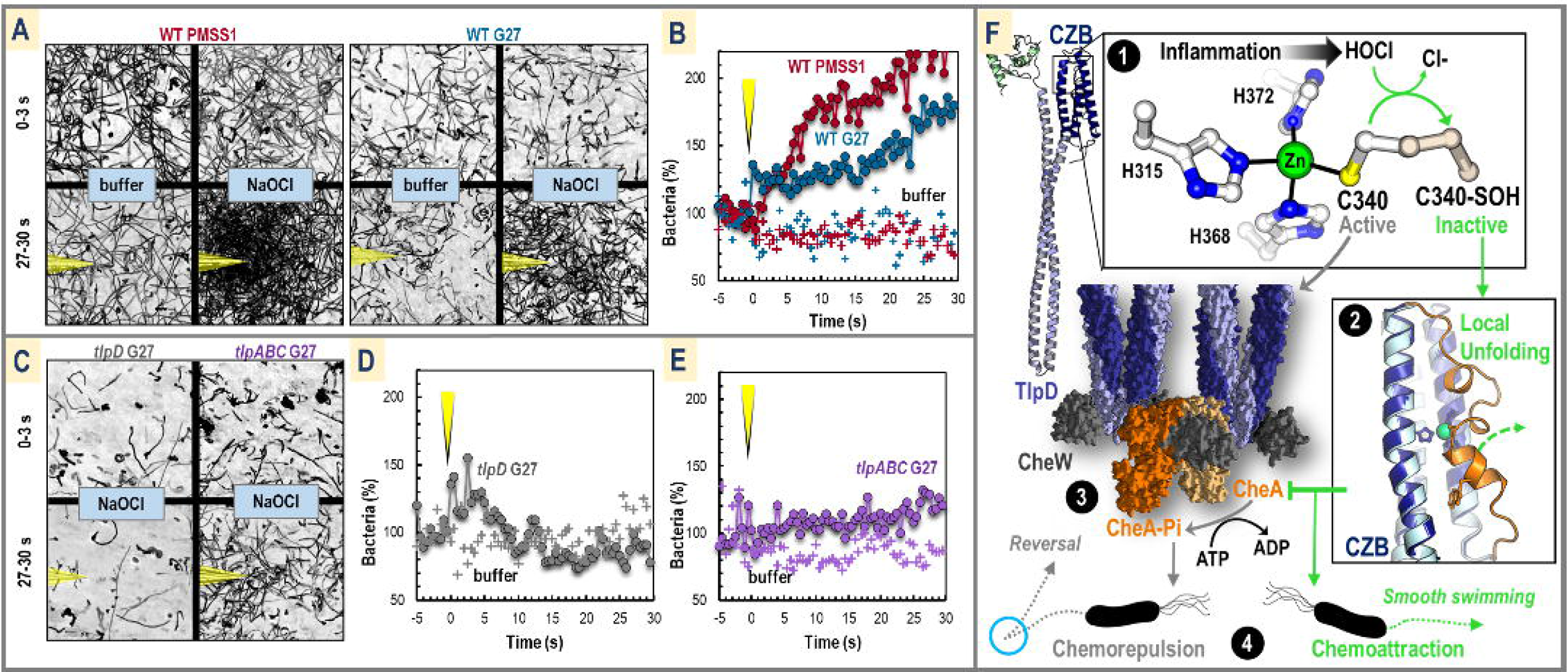
*H. pylori* Exhibits TlpD-Dependent Chemoattraction to Sources of Bleach. (A) Data from point-source chemotaxis experiments (Huang et al., 2015) are shown for two different *H. pylori* strains using a highly-calibrated micropipette for chemoeffector delivery. Three second motility traces are shown prior to treatment (0-3 s) and following 30 s of treatment (27-30 s) with buffer (500 mM sodium phosphate, pH 6.7) or 10 mM NaOCl (diluted into buffer, pH 6.7). For both strains PMSS1 (red) and G27 (blue) attraction is seen for NaOCl with a large increase in the number of bacteria appearing in the field of view following 30 s of treatment. The position of the micropipette is highlighted in yellow in the treatment panels. (B) Data from A is quantified over time with the average percentage of bacteria in the field of view five seconds prior to treatment normalized to be 100 % (−5-0 s). Treatment with buffer (pluses) or 10 mM buffered NaOCl (circles linked by lines) begins at time zero, noted with the yellow micropipette. Each data point shown is the average over 2 s of video (60 frames). (C) Experiments performed as in A for the *tlpD* G27 (gray) and *tlpABC* (purple) G27 mutants with NaOCl. (D) Quantified data for *tlpD* shows TlpD is required for chemoattraction to NaOCl. (E) Quantified data for *tlpABC* experiments shows a weaker attraction can be retained with only TlpD, increasing the bacteria in the field of view by approximately 25 % after 30 s of treatment. See Fig. S4 for videos of these experiments and Fig. S4D for statistical analysis. (F) Shown is a synthesis of the data from this work as the basis for a new model for how TlpD functions to sense bleach as a chemoattractant. By default, TlpD is active and promotes swimming reversals (blue circle) by stimulating CheA autophosphorylation (gray arrows). TlpD is deactivated by HOCl-sensing, leading to smooth swimming and chemoattraction (green arrows). The inflammation product HOCl oxidizes a specially-tuned C340 to form a cysteine sulfenate (panel 1). Oxidation of C340 causes the Cys to detach from the Zn-binding core, and a large conformational change occurs in the receptor (panel 2). This structural rearrangement deactivates the receptor so that it no longer stimulates CheA autophosphorylation in the chemotaxis signaling complex (panel 3). Decreasing the cellular pool of CheA-Pi leads to increased smooth swimming and chemoattraction to the HOCl source.

## DISCUSSION

We report the unexpected discovery that the prevalent gastric pathogen *H. pylori*, a causative agent of stomach cancer, is attracted to sources of bleach by virtue of its cytosolic chemoreceptor TlpD. By reconstituting the chemotaxis complex *in vitro* we determined TlpD contains a conserved C340 required for promoting the autophosphorylation of CheA (Fig. 1-3). Further biochemical analyses revealed C340 is tuned to be highly and preferentially reactive toward HOCl and that oxidation of C340 by HOCl stimulates a large conformational change in the receptor that presumably inhibits CheA activity, leading to a decrease in the cellular pool of CheA-Pi (Fig. 4). TlpD regulation of CheA activity shows half-maximal inhibition at 100 μM HOCl, and our data suggest this sensing is strongly cooperative at the level of the TlpD homodimer (Fig. 4), consistent with the array performing signal amplification as is seen for the *E. coli* chemotaxis system (Hazelbauer et al., 2008). We expect that for *H. pylori,* oxidation of just a few TlpD receptors *in vivo* can inactivate much of the available CheA kinase and thus the cytosol need not reach HOCl concentrations high enough to oxidize all of the TlpD pool to elicit an attraction response. Experiments with chemotaxing *H. pylori* show exposure to bleach at concentrations as low as 250 μM decreases reversals and promotes smooth swimming (Fig. 5) and the bacteria rapidly localizes toward a point-source of bleach (Fig. 6). Bleach attraction was TlpD-dependent and retained in two *H. pylori* strains isolated from distant geographic locations (Fig. 6).

### Cys340 Participates in a Cys-Zn Switch to be Highly and Preferentially Reactive with HOCl

The molecular basis for TlpD’s specific responsiveness to bleach is the preferential reactivity of Cys340 with HOCl versus the chemically-similar H_2_O_2_. Such specificity can be understood based on differences between the reactivity of these species with protein residues. H_2_O_2_ in the micromolar range almost exclusively reacts with cysteine, and to a lesser extent with methionine, whereas the targets of HOCl are more broad and exhibit extremely fast reaction rates; a detailed kinetic analysis measured the reaction rate of HOCl with Cys to be 3.0 × 10^7^ M^−1^ s^−1^, and 3.8 × 10^7^ M^−1^ s^−1^ for Met (Lebrun et al., 2016). That same study showed cysteines bound to zinc are by far the most reactive protein residue with HOCl, achieving a near diffusion-limited rate of 9.3 × 10^8^ M^−1^ s^−1^, almost 3 orders of magnitude greater than cysteine alone (Lebrun et al., 2016), and in fact also exhibit decreased reactivity with hydrogen peroxide and oxygen (Bourlès et al., 2011). Previous work has identified numerous other proteins containing zinc-bound cysteines that are targets for HOCl oxidation including Hsp33 (Lebrun et al., 2016), and alcohol dehydrogenase, which is 3000-fold more reactive with HOCl than peroxide (Crow et al., 1995). The example we found to be most similar to TlpD is the activation of human neutrophil collagenase that occurs by HOCl oxidation of a single zinc-bound Cys, a mechanism coined as a “Cys-Zn switch” (Springman et al., 1990; Van Wart and Birkedal-Hansen, 1990). Together, our data indicate the molecular function of TlpD is modulated by reaction of HOCl with C340 to switch between two conformations: (1) a “reduced” and Cys-zinc-bound state that is fully folded and activates CheA, and (2) an “oxidized” cysteine sulfenate state with a partially unfolded CZB domain that does not activate CheA, and that can be reduced and resurrected by cellular reductants (Fig. 6F).

### Bleach is a Prevalent Host-Produced Oxidant Encountered by Helicobacter pylori

HOCl, which is produced by the granulocyte-specific enzyme myeloperoxidase from H_2_O_2_ and chloride anion, reaches levels of up to 5 mM at sites of inflammation (Pullar et al., 2001; Weiss, 1989). In contrast, maximal gastric concentrations of extracellular H_2_O_2_ are thought to be 1-15 μM and perhaps rarely reach ~100 μM in the presence of oxidants from dietary sources such as coffee (Halliwell et al., 2000; Schröder and Eaton, 2008). The concentration of superoxide is approximately 2 μM and its half-life is on the order of milliseconds and decomposes rapidly (Ganesana et al., 2012). Previous analyses of *H. pylori* oxidant sensing (Behrens et al., 2016; Collins et al., 2016) used millimolar H_2_O_2_ and superoxide, which are much higher concentration than what occurs *in vivo*. Additionally, although triggered human neutrophils produce several ROS during inflammation, including H_2_O_2_ and superoxide, it seems these are all funneled into the production of HOCl as the major product, as quantitative analyses show essentially all H_2_O_2_ generated is converted to HOCl (Test and Weiss; Weiss, 1989).

Bleach is a potent disinfectant that kills bacteria at micromolar concentrations and is sufficiently cell-permeable to oxidize intracellular thiols at sub-lethal doses (Tatsumi and Fliss, 1994). We showed that *H. pylori* is exceptionally resistant to bleach, likely due to its strong antioxidant defenses including three peroxiredoxins, and high expression of its catalase, KatA (Stent et al., 2011). KatA in particular has been shown to be key for the bacteria to survive ROS bursts from triggered immune cells, and evidence indicates this resilience leads to the chronic release of ROS that damages gastric tissue and results in disease (Ramarao et al., 2000).

Our finding that *H. pylori* is attracted to sources of HOCl by TlpD suggests that *H. pylori* does not just tolerate, but can actively seek out sources of HOCl, which would be produced by neutrophils at sites of inflammation. Neutrophils are a hallmark of *H. pylori* infection and are actively recruited in response to *H. pylori* induction of interleukin-8 (Dixon et al., 1996). Neutrophils concentrate near the stem precursor and cell zones of the gastric gland epithelium (Lee, 2014) which undergoes proliferation in response to *H. pylori* gastric gland colonization (Sigal et al., 2015). *H. pylori* colonization of the gastric glands is an important strategy for the bacteria’s persistent infection of the stomach and requires chemotaxis (Howitt et al., 2011; Keilberg et al., 2016), and therefore chemoattraction to HOCl may constitute a strategy *H. pylori* uses to persist in neutrophil-infiltrated gastric glands.

This behavior is consistent with other pro-inflammatory strategies of *H. pylori,* such as its degradation of the host-secreted extracellular anti-oxidant glutathione (Suzuki et al., 2012), and that its colonization levels are reduced in mice treated with the ROS scavenger ursodeoxycholic acid (Thao et al., 2008) or sulforaphane, which stimulates host antioxidant protein production through Nrf2 (Yanaka et al., 2009). In this regard, *H. pylori* belongs to a growing list of bacterial pathogens and pathobionts that can promote and benefit from host inflammation, thereby setting up positive feedback loops that can drive chronic inflammation (Litvak et al., 2017; Paiva and Bozza, 2014). For example, *Salmonella*-induced inflammation increases the concentrations of colonic tetrathionate and nitrate, which it recognizes as chemoattractants, and it can use these inflammation products as electron acceptors in anaerobic respiration to outcompete fermentation-restricted members of the native microbiota (Bäumler and Sperandio, 2016).

### H. pylori Integrates Multiple Chemical Cues to Persist in the Gastric Environment it Shapes

*H. pylori* is a notoriously persistent colonizer often retained for an individual’s lifetime, causing chronic gastritis that can proceed to ulcers or adenocarcinoma. *H. pylori* requires chemotaxis to colonize the stomach and seek out injured gastric tissue (Aihara et al., 2014; Huang et al., 2017; Rolig et al., 2012). To persistently colonize the stomach, *H. pylori* uses a limited chemoreceptor repertoire to integrate information about chemical gradients, both pre-existing and of its own making. These receptors often sense multiple chemoeffectors and likely form intermingled arrays (Collins et al., 2016), raising the possibility that receptors integrate information for chemoeffectors to which they do not directly bind. Interestingly, many of these chemoeffectors reflect the environment of the *H. pylori*-infected, as opposed to a naïve stomach, such as low concentrations of urea (Huang et al., 2015) and presence of autoinducer-2 (Anderson et al., 2015), consistent with a role for chemotaxis in persisting within the chronically infected stomach. Now, we add to this picture chemoattraction to HOCl produced by *H. pylori*-induced neutrophil infiltration. The fact that TlpD mediates both direct HOCl sensing and pH responses complicates what conclusions can be drawn from *tlpD* mutants (Collins et al., 2018; Huang et al., 2017; Rolig et al., 2012; Schweinitzer et al., 2008) however consistent with a role in TlpD mediating chemoattraction toward HOCl, *H. pylori* deficient in *tlpD* exhibit defects in stomach colonization and fail to persist in the antrum, the region of highest inflammation (Huang et al., 2017; Rolig et al., 2012). In conclusion, our findings reveal that *H. pylori* has the surprising capacity of chemoattraction toward bleach, a compound commonly viewed as a noxious oxidant and antimicrobial, which may help enable its persistence in chronically inflamed host tissue. More generally, CZB-domain containing proteins are encoded by the genomes of numerous host-associated bacterial species (Draper et al. 2011a), suggesting the broad utility of HOCl-sensing for bacteria living together with animals that produce bleach as an antimicrobial response.

## ACKNOWLEDGEMENTS

We acknowledge our colleagues at University of Oregon, especially Dr. Daniel Shoup, the lab of Dr. Alice Barkan, Dr. Pete Von Hippel, and Dr. Steven Weitzel for use of laboratory space, equipment, and training. We also thank Dr. Karen Ottemann at UC Santa Cruz who participated in helpful discussions regarding our data and very generously provided the *H. pylori* strains used in this work.

## AUTHOR CONTRIBUTIONS

AP designed and performed most of the experiments and analysis, wrote the manuscript, and created the figures. DT purified some of the proteins used, performed fluorescence experiments with the C340A mutant, and contributed to data collection and processing of motility velocity assays. MRA performed the microgradient point-source chemotaxis assays. SJR and KJG contributed to conceptualizing and designing experiments, data analysis, and provided critical review and editing of the manuscript.

## STAR METHODS

### Contact for Reagent and Resource Sharing

Further information and requests for resources and reagents should be directed to and will be fulfilled by the Lead Contact, Dr. Arden Perkins.

### Experimental Model and Subject Details

#### Helicobacter pylori Growth and Preparation

The high motility strain *H. pylori* G27 was used for most *in vivo* assays and all mutants were created in this background previously (Collins et al., 2016) and supplied as a gift from Dr. Karen Ottemann (UC Santa Cruz). As an additional control, strain PMSS1 was used in microgradient point-source assays. For growth, frozen stocks were used to inoculate blood plates containing 5-percent defribrinated horse blood (Hemostat Laboratories), 4.4-percent w/v columbia agar, 0.02 mg/ml β-cyclodextrin, 0.016 mg/ml amphotericin B, and 0.02 mg/ml vancomycin. Bacteria were grown on plates for 72 hours at 37° C, 10 % CO_2_, before passaging to a fresh blood plate and allowed to grow an additional 24 hours. For motility assays, scrapes from blood plates were used to inoculate 3 ml of brucella broth supplemented with 10 % fetal bovine serum and 10 μg/ml vancomycin (BB_10_) for 6 hours at 37° C and 10 % CO_2_ shaking at 200 rpm. The liquid cultures were then removed from the shaker and allowed to incubate without shaking for another 1-3 hours to obtain maximum motility. Prior to motility and chemotaxis assays, the cultures were diluted with fresh BB_10_ to approximately 0.1 O.D.

#### Escherichia coli Growth and Preparation

*E. coli* strain UU1250, which is engineered to lack all chemoreceptors and is commonly used in chemotaxis studies (Liu and Parales, 2008), was supplied as a gift from Dr. Sandy Parkinson (University of Utah). Frozen stocks of *E. coli* were used to inoculate 25 ml of sterile LB media and was grown shaking overnight at 37 ° C. In the morning, 100 μL of the overnight culture was diluted into 5 ml of fresh LB media and allowed to grow until reaching an O.D. of approximately 0.5, and then was used for motility assays.

### Method Details

#### Sequence Analysis and Conservation of TlpD

The full-length protein sequence of *Helicobacter pylori* TlpD from strain SS1 (Uniprot entry A0A1U9IUC7) was used for BLAST (Altschul et al., 1990) searches of the non-redundant sequence database using default threshold values. We performed one search for only *H. pylori* sequences and a second search excluding *H. pylori* sequences. We manually curated these sequences to retain only those that were chemoreceptors and also contained a C-terminal CZB domain. Of these, some sequences lacked the N-terminal region corresponding to approximately the first 120 residues; because we were interested in analyzing the conservation of TlpD cysteine positions, and three of those positions were contained in the N-terminal region, we imposed an additional restriction that we only included sequences in the analysis that lacked no more than the first 30 N-terminal residues. This resulted in a total of 459 putative TlpD homologues. We performed a multi-sequence alignment of this subset using Clustal Omega (Sievers et al., 2011) and generated a relatedness tree using PhyML (Guindon et al., 2010) (Fig. 1C).

To create the homology model of TlpD, we submitted the *Hp*TlpD (SS1 strain) sequence to the i-Tasser server (Zhang, 2008) as two separate pieces corresponding to the N-terminal region (residues 1-120) and the coiled-coil and CZB domain (residues 121-433), and the subsequent predicted models were manually joined (Fig. 1D). While generation of the model could be guided by crystal structures of the chemoreceptor coiled-coil region and a CZB domain from *E. coli* DgcZ, the TlpD N-terminal region is unlike any experimentally-determined structure and so the fold of this region is remains highly speculative. We performed a sequence conservation analysis at the five cysteine positions found in *Hp*TlpD (C35, C103, C117, C308, C340) for *H. pylori* sequences and *non-pylori* sequences using the WebLogo server (Crooks, 2004) and mapped these onto the i-Tasser homology model (Fig. 1D-E).

#### Purification of Recombinant Proteins

pBH plasmids with ampicillin resistance and a T7 promotor system (Goers Sweeney et al., 2012) were ordered from GeneScript that contained the sequences for *Helicobacter pylori* TlpD (SS1 strain, Uniprot: A0A1U9IUC7) and C340A mutant, CheW (SS1 strain, Uniprot: A0A1U9ITT8), CheA (G27 strain, Uniprot: B5Z859), and *Salmonella enterica* typhimurium McpA (LT2 strain, Uniprot: Q8ZM22) each with the N-terminal His-tag: MGHHHHHHDYDIPTTENLYFQGS. For these constructs, ArcticExpress DE3 competent *E. coli* cells (Agilent) were transformed with plasmids by heat shock and plated on +ampicillin LB agar plates. After overnight growth at 37° C, single colonies were selected for growth and large scale protein expression. Briefly, individual colonies were used to inoculate 25 ml LB/+AMP cultures and grown overnight shaking at 37 °C. The following morning 5 ml of overnight culture was added to 4×1 L cultures of LB/+Amp, and allowed to grow shaking at 37° C to O.D. 0.6-0.8. Cultures were then transferred to a temperature-controlled incubator and allowed to grow shaking for 30 minutes at 10° C, prior to induction with 1 mM IPTG and growth overnight at 10° C. After approximately 16 H, cultures were harvested by centrifugation at 5k rpm at 4° C. Similar protocols were used for expression of the CZB domain of *E. coli* DgcZ (DgcZ-CZB in strain K12), which was supplied as a gift from Dr. Tilman Schirmer of University of Basel (Zähringer et al., 2013), using expression with Rosetta cells and a pet19b vector.

For protein purification, cells were diluted into ice-cold lysis buffer containing 10 mM imidazole, 50 mM HEPES, 10 % glycerol, 300 mM NaCl, 0.5 mM TCEP, pH 7.9. Cells were kept on ice and lysed by sonication and then centrifuged at 15k rpm to separate out the insoluble fraction. The soluble portion was retained and applied to a pre-packed gravity column of Ni-NTA agarose beads (Qiagen) equilibrated with lysis buffer. Lysate was incubated with the beads for 30-60 mins and allowed to flow through the column over the beads twice. The column was then washed with lysis buffer to remove contaminates until no further protein was observed in the flow through, as monitored using a Bradford assay. Protein was eluted by the addition of lysis buffer containing 300 mM imidazole. Elution fractions were judged for purity by SDS-PAGE and pooled before further purification by gel filtration on an Akta FPLC. All proteins were obtained in high yields on the order of 40-100 mg per prep, and fractions containing pure protein were dialyzed into appropriate buffers and concentrated; TlpD, TlpD-C340A, and CheA were found to be highly soluble and stable up to 300 μM, but CheW could only be concentrated to approximately 150 μM without precipitation. Proteins were flash frozen in liquid nitrogen and stored at −80° C.

#### Preparation of Bleach Solutions

High grade sodium hypochlorite (NaOCl) was obtained from Sigma-Aldrich as a 10-15 % solution. Precise concentration of NaOCl was assayed directly using the extinction coefficient of 350 M^−1^ cm ^−1^ measured at 293 nm in water (Furman and Margerum, 1998). Bleach stock was kept cold and in the dark until needed, and bleach solutions for assays were made fresh. Stock solutions of NaOCl are extremely basic, and so in all cases the pH of bleach solutions was measured and adjusted as needed prior to addition, and great care was taken to ensure the final pH of reactions was not perturbed. The pKa of bleach is 7.53, and so the experiments performed here contain a mixture of hypochlorous acid (HOCl) and hypochlorite (^−^OCl) in solution, but for simplicity we refer to these solutions simply as either HOCl or NaOCl.

#### ATP Radio-labeling and Functional Assays of Reconstituted Signaling Complex

Unless otherwise stated, all functional assays were performed at 20° C in standard kinase buffer containing 50 mM Tris pH 7.5, 100 mM NaCl, and 10 mM MgCl_2_. Typical reactions contained purified chemotaxis proteins in approximately stoichiometric concentrations: 4 μM CheA, 8 μM CheW, and 24 μM TlpD dialyzed into kinase buffer. Treatments of bleach, peroxide, paraquat, TCEP, and zinc acetate were added to proteins 1 H prior to the start of the reaction. ATP solutions were prepared with a 500:1 ratio of cold ATP (Thermofisher) to hot ATP [◻-32P] (Perkin Elmer) diluted into kinase buffer. Time courses were carried out in a 96-well clear bottom plate (Corning) with reactions started by the addition of ATP. At each timepoint, 3-10 μl samples were extracted and immediately quenched in a solution of 190-197 μL of 75 mM H_3_PO_4_ and 1 M NaCl. Subsequently, samples were added to a Whatman Minifold 96-well slot blotter and drawn through a HyBond PVDF nitrocellulose membrane by vacuum. Three additional washes with 200 μL quenching solution were performed to remove excess ATP so that only bound protein remained. The nitrocellulose was dried and then imaged using a phosphoimaging cassette (Molecular Dynamics) with a 30-60 minute exposure, and then scanned using a Storm 825 imager (GE Healthcare). Data was quantified by densitometry using Image Studio. Identical-sized circles were used for all measurements, and raw values from experimental samples were normalized against “time zero” samples that contained only ATP and no kinase.

We performed similar experiments using wild type TlpD, CheW, and CheA across a variety of pHs to test if TlpD may be involved in directly sensing pH. TlpD has been implicated to be involved in chemorepulsion from acid and chemoattraction to alkaline pH (Huang et al., 2017), and if this sensing was direct we expect in the reconstituted chemotaxis complex assay this would be reflected as higher CheA activity at more acidic pH, leading to more swimming reversals and chemorepulsion, and less CheA activity at basic pH. Our results actually found the opposite to be true, as CheA autosphosphorylation activity increased at more alkaline pH (Fig. S1). This has been previously reported for studies with *E. coli* CheA alone (Hess et al., 1988), suggesting that the increase in activity may be due to the CheA autophosphorylation reaction being more favored at alkaline pH, although further work is needed to confirm this. However, activity is stable near neutral pH, and because TlpD is within the well-buffered cytosol it is unclear if the changes in activity we observe at extreme pHs have physiological relevance for signaling. It remains possible TlpD could sense pH through binding a titratable buffer component, but no evidence supporting such a mechanism has been reported.

#### Detection of Cysteine Sulfenic Acid by Western Blotting

Experiments were prepared with either 20 μM TlpD, TlpD-C340A, *E. coli* DgcZ-CZB, or *Salmonella* McpA in PBS buffer pH 7 with 10 mM freshly prepared dimedone (Sigma). The proteins were treated with various concentrations of NaOCl for 1 H at 30 ° C and then 5 μl of sample was quenched into 195 μl of quenching buffer (see above) and drawn through a HyBond PVDF nitrocellulose membrane by vacuum. The membrane was washed three times with quenching buffer and then once with TBST buffer (50 mM Tris pH 7.5, 150 mM NaCl, 0.1 % Tween-20). Rabbit anti-cysteine sulfenic acid primary antibody from Kerafast was used to detect dimedone-modified cysteines (Seo and Carroll, 2009) using a 1:4000 dilution in a blocking buffer of 5 % milk in TBST. The membrane was incubated with primary antibody overnight with gentle rocking. The following day the membrane was washed three times with 20 ml of TBST for 15 minutes, followed by incubation with anti-rabbit-HRP secondary antibody (Santa Cruz Biotechnology) at a 1:4000 dilution in blocking buffer for 1 H. The membrane was again washed three times with 20 ml of TBST for 15 minutes, and then a Pierce ECL detection kit was used to visualize the blot using chemiluminescence.

#### Mass Spectrometry

20 μM of pure recombinant TlpD in PBS buffer pH 7 was prepared for analysis by treatment with 1 mM TCEP, H_2_O_2_, or NaOCl for 1 H. Subsequently, the samples were alkylated by the addition of 5 mM iodoacetamide and allowed to react for 30 minutes at room temperature sheltered from light. The sample was then buffer-exchanged with fresh buffer three times by centrifugation using a 10 kDa cutoff and flash frozen in liquid nitrogen prior to analysis. Mass spectrometric analysis of TlpD was performed as a service by Dr. Larry David at Oregon Health and Science University’s (OHSU) Proteomics Shared Resource Facility. As per their request, we include the following statement: “Mass spectrometric analysis was performed by the OHSU Proteomics Shared Resource with partial support from NIH core grants P30EY010572, P30CA069533, and S10OD012246.” Protein samples were subjected to digestion with LysC, and 3 μg of digest was analyzed on a Q-Exactive HF instrument using a 75 μm × 250 mm nanospray C18 column. MS/MS data were searched with Sequest against an *E. coli* database supplemented with the TlpD protein sequence. False discovery was controlled using a reversed sequence database and only q-scores below 0.05 were accepted. C340 was located within the peptide NCRLGKWYYEGAGK (567.94415 Da), and carbamidomethylated (+57.02146 Da), sulfinate (+31.98983 Da), and sulfonate (+47.98474 Da) forms were detected. Ion extractions were performed for the +3 charge states for the three forms of this peptide using a 2 ppm tolerance, and peaks were integrated to obtain absolute values for quantification and comparison of modifications across experiments (Fig. 4E, Fig. S2A-C).

#### Fluorescence, CD Spectroscopy, and Analytical Ultracentrifugation

For Zn-chelation assays, we used the fluorescent probe Zinpyr-1 (Abcam) with excitation at 492 nm and maximum emission near 527 nm. 5 mM stock solutions of Zinpyr-1 were prepared in DMSO and diluted to 50 μM final concentration for experiments with zinc controls and TlpD in a buffer of 100 mM Tris pH 7, 300 mM NaCl. Samples were loaded into a 2×2 mm quartz cuvette (Starna Cells Inc.) and fluorescence was monitored at 20° C using a FluoroMax-3 Spectrofluorometer (HORIBA Scientific). For time courses, fluorescence was measured every 60 s over the course of 3 H. Identical samples containing only buffer were used as blanks, and final spectra shown are blank-subtracted spectra, but these were minimally different (Fig. 3A-B). For intrinsic protein fluorescence assays with TlpD and the C340A mutant we used an excitation wavelength of 295 nm, to optimize signal from the single Trp345, and monitored emission in the 300-400 nm range. Samples were prepared in a buffer of 25 mM NaCl and 20 mM Tris pH 7 and experiments were conducted at 20° C. Samples were pre-treated for 1 H prior to measuring fluorescence for endpoint fluorescence spectra of TlpD bleach treatments (Fig. 4G-I), or subsequently treated with TCEP for 30 minutes (Fig. 4J). Fluorescence anisotropy experiments for recombinant TlpD in PBS buffer pH 7 and 1 mM TCEP were collected with excitation at 295 nm and monitoring emission at 340 nm. Intrinsic fluorescence was too weak at low protein concentrations to be reliably detected and so a K_D_ value could not be calculated from this data, but were still useful to indicate that the protein is fully oligomerized by approximately 16 μM (Fig. 2B)

Protein CD was performed for TlpD with a Jasco J-810 spectropolarimeter using a 1 mm pathlength quartz cuvette at 20° C. Samples were prepared with 20 μM protein in a buffer of 25 mM NaCl and 20 mM Tris pH 7 and addition of various concentrations of buffered NaOCl (Fig. S2E). CD is sensitive to electrical conductivity from salt and signal-to-noise ratios become worse as the photomultiplier tube voltage increases (Greenfield, 2006), and in our samples addition of millimolar quantities of NaOCl resulted in spectra deterioration. Because the overall shape of the curves of untreated protein and samples treated with millimolar NaOCl are highly similar, we expect that the observed decrease in signal is due to high photomultiplier tube voltages (>500 V) and not to global protein unfolding (Fig. S2E). All spectra for fluorescence and CD experiments were corrected for background by subtracting signal from buffer-only samples.

Analytical ultracentrifugation experiments were performed for recombinant TlpD using a Beckman Proteome Lab XL-I centrifuge and a 60 Ti four-cell rotor. Interference experiments using various concentrations of TlpD in PBS pH 7 and 1 mM TCEP with a total volume of 400 μl were performed at 55,000 rpm. Data were processed with SedFit (Lebowitz et al., 2002) using a continuous distribution model with rmsd of the final datasets < 0.0078 (Fig. 2B).

#### Bacterial Motility Velocity Assays

Bacteria were grown as described above and diluted in fresh media to be approximately O.D. 0.1. For treatments, 2 μl of motile bacteria combined with 2 μL of chemotaxis buffer (10 mM PBS pH 7, 1 mM EDTA) or bleach diluted into chemotaxis buffer from a concentrated stock of bleach pH’d to 7. The samples were mixed by gentle pipetting, applied to a 10-well slide (MP Biomedicals) and covered with a 22 mm micro cover glass (VWR), and visualized immediately. For each experiment, brightfield videos of swimming bacteria were recorded using a Nikon Eclipse Ti inverted scope using a 20x objective and equipped with an AirTherm temperature-controlled sample chamber set to 37 ° C. Videos were 30 s in duration and recorded at 25 frames per second (Fig. 5A). The number of bacteria tracks counted for each bleach concentration were as follows: *E. coli:* buffer (409), 500 μM (377), 1 mM (283), 2 mM (324), 3 mM (no motile bacteria detected); *H. pylori* buffer (91), 500 μM (39), 1 mM (33), 5 mM (70), 6 mM (60), 7 mM (23), 8 mM (no motile bacteria detected).

#### Bacterial Chemotaxis Assays

*H. pylori* were grown as described above and diluted in BB_10_ to be approximately O.D. 0.1 and checked for motility after 30-60 minutes; the bacteria were used for motility and chemotaxis assays once the culture was observed to be near 100 % motile. For *H. pylori* swimming assays, we constructed small chambers that allowed for the rapid delivery and visualizing of chemoeffector treatments. Two parallel strips of double-sided tape spaced 5 mm apart were placed lengthwise along a 22×60 mm microscope cover glass (Fisher Scientific) and a 24×40 mm microscope cover glass (VWR) was placed on top and sealed with gentle pressure. The resulting chamber is approximately 5 mm × 40 mm × 0.1 mm and holds approximately 20 μL of volume. To record brightfield videos of motile *H. pylori* G27 wild type and mutants we used a Nikon Eclipse Ti inverted scope using a 40x objective equipped with an AirTherm temperature-controlled sample chamber set to 37 ° C. For chemoeffector treatments, 10 μl of cells were added to the chamber and a 30 s video was recorded to document baseline swimming, and then 10 μl of 2x treatment solution was flowed into the chamber and a 3 min video was recorded once the initial flow subsided (< 10 s after treatment). Though this method effectively dilutes the bacterial O.D. by half, no major differences in swimming behavior were observed between untreated and media or buffer-treated bacteria (see Fig. S3F). All treatments were prepared in identical proportions: chemoeffector (urea, HCl, or NaOCl) was diluted into 200 μL of 100 mM Tris pH 7, and then 800 μl of BB_10_ that was pre-incubated with 10 % CO_2_ for 30 min was added, for a total volume of 1 ml. NaOCl treatments were prepared fresh and applied to bacteria immediately after addition to BB_10_ (Fig. 5C-E).

To test if HOCl-induced swimming changes are TlpD-dependent, assays were performed with a *tlpD* mutant, however previous studies have shown *H. pylori tlpD* mutants adopt a mostly smooth-swimming baseline behavior (Behrens et al., 2016; Collins et al., 2016), and thus increases in smooth-swimming cannot be observed relative to swimming in media or buffer alone. There is some evidence showing *H. pylori* chemoreceptors form intermingled arrays at cell poles (Behrens et al., 2016; Collins et al., 2016), and a shift to smooth swimming was also seen for a *tlpB* mutant (Goers Sweeney et al., 2012), so it may be that deleting a chemoreceptor partially disrupts array formation *in vivo*. We therefore performed assays with the *tlpD* mutant in a background of BB_10_ with the addition of 50 mM HCl, which can be sensed through TlpA and TlpB, to promote enough reversals to be useful for comparisons with HOCl treatments. While *tlpD* in BB_10_ alone showed ~100 % smooth-swimming, addition of 50 mM HCl dropped this to approximately 30 %, comparable to wild type without HCl addition (Fig. S3G). Cotreatment of the *tlpD* strain with 100 mM HCl and 1 mM HOCl showed a negligible increase of smooth-swimming of 5 % over HCl treatment alone, supporting that HOCl-sensing is TlpD dependent (Fig. 5E and Fig. S3G).

Point-source micropipette assays were performed as previously described (Huang et al., 2015). An Eppendorf Femtotip II microinjection micropipette, loaded with either a buffer of 500 mM sodium phosphate pH 7 or 10 mM NaOCl diluted into buffer, was inserted into a swarm of motile *Helicobacter pylori* using an Eppendorf micromanipulator. The pH of treatment solutions was measured prior to experiments to ensure addition of NaOCl did not shift them to be alkaline. Treatment was initiated by applying a pressure of 30 hPa to maintain a constant flow of 0.372 pl per minute, which is low enough to not physically perturb the swimming bacteria (Huang et al., 2015). Videos of swimming bacteria were captured at 30 frames per second on a Zeiss Axiovert-35 inverted microscope and experiment temperature was kept stable at 37 ° C with a heated stage (Fig. 6A-E, Fig. S4). For each experiment, a baseline for the number of bacteria in the field of view was established pretreatment, and then the micropipette was lowered into the swarm and a gentle flow of either sodium phosphate buffer or 10 mM NaOCl diluted into buffer (pH 7) was initiated.

### Quantification and Statistical Analysis

#### Curve fitting of reactions and calculation of thermodynamic constants

Radio-ATP time course assays were run to completion (15 min-2 H depending on the sample) and raw data was normalized with the assumption that endpoint values reflected maximally phosphorylation CheA. For Michaelis-Menton assays using CheA and varying [ATP], the data were fit assuming the second order reaction proceeds as: CheA + ATP ↔ CheA-Pi + ADP; so that the radioactivity measured is directly proportional to [CheA-Pi], and [CheA] and [ATP] can be deduced for any timepoint by knowing initial concentrations. Thus, experimental data were fit to the second-order rate equation:

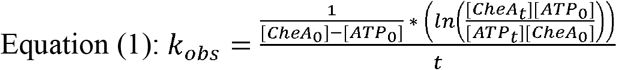

Where [CheA_0_] is the starting concentration of CheA, [ATP_0_] is the starting concentration of ATP, [CheA_t_] is the concentration of CheA measured at time *t*, and [ATP_t_] is the concentration of ATP measured at time *t*. A least-squares fit of the mean data values to Eq. 1 were performed using Microsoft Excel’s Solver addon to minimize the error of the fit with the data values weighted by the standard deviation of each sample, calculated as:

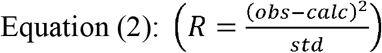

Where R is error, *obs* is the measured value of [CheA] and *calc* is the calculated concentration of [CheA] based on Eq. (1), and *std* is the standard deviation of the sample measurements. Individual k_obs_ values from time courses were plotted as a function of ATP concentration, with error calculated as in Eq. 2, and fit to the Michaelis-Menton equation (Fig. 2A):

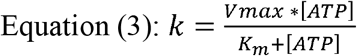

Where *k* is the reaction rate, *V*_*max*_ is maximum velocity, *K*_*m*_ is the substrate concentration required for half-maximal velocity, and [ATP] is the concentration of ATP. This resulted in a calculated K_m_ of CheA for ATP of 136 μM, and V_max_ of 0.0033 μM^−1^ min^−1^, indicating that CheA autophosphorylation is extremely slow in the absence of activators. Subsequent time course reactions were run under pseudo first-order conditions using 1 mM ATP and k_obs_ values were calculated from fits of the data as described above to the pseudo-first order rate equation:

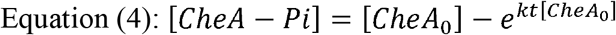

For determining kinetically-defined K_D_ values for signaling complex components (Fig. 2D-E) we performed a series of time course reactions titrating a single protein component, either CheW or TlpD, against 4 μM CheA and 1 mM ATP, and all other components were kept at a constant concentration. The data for each time course was fit to Eq. 4 to calculate k_obs_, and the series of k_obs_ values were normalized assuming maximal values corresponded to fully-bound complex (either CheA-CheW, or CheA-CheW-TlpD). The k_obs_ values were plotted against concentration of the titrated component and fit to a two-component binding (Hill) equation:

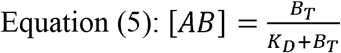

Where it is assumed binding occurs as A+B↔AB, [AB] is the concentration of the bound complex, B_T_ is the total added titrated component, and K_D_ is the thermodynamic dissociation constant. Using this method, we calculated the K_D_ of complex formation of CheA+CheW↔CheA-CheW to be 14.6 μM. In the case of the titration of TlpD against CheA and CheW (Fig. 2E) we were unable to use a saturating amount of CheW due to the limits of the protein binding capacity of the nitrocellulose membrane. Instead, we used 4 μM CheA and 40 μM CheW and approximated the concentration of the CheA-CheW complex to be 2 μM, based on our previous titration of CheW and CheA, although this equilibrium could be shifted by TlpD addition. However, data were well-fit using this estimate, producing a K_D_ for TlpD to CheA-CheW of 15.2 μM, and test fits using values between 2-4 μM CheA-CheW complex only subtly changed the calculated K_D_ by ~1 μM, so we are fairly confident in this calculation despite that we did not directly measure the CheA-CheW complex concentration. Equation 5 was also used to fit quantified data from fluorescence and western blot experiments using Hill coefficients of 1 and 2 (Fig. 4D,H).

Calculation of the TlpD dimerization K_D_ was performed by integrating the monomer and dimer peaks from analytical ultracentrifugation experiments to obtain the monomer/dimer ratio, which directly relates to the K_D_ of a two-species oligomerization process by the equation:

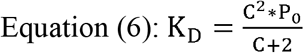

Where C is the ratio of monomer to dimer and P_0_ is the total protein concentration (Fig. 2B).

#### Tracking of Bacterial Swimming

For motility velocity assays we used in-house MATLAB-based automated particle tracking software from the laboratory of Dr. Raghuveer Pathasarathy (University of Oregon) to track bacteria swimming paths (Parthasarathy, 2012). Videos of bacteria were first converted to a series of tif files using ImageJ (Collins, 2007), and inverted so that bacteria appeared as bright spots on a dark background, and then input into the tracking software. Spots of intensity greater than 0.998 standard deviations above the mean were defined as “objects” and nearest neighbors between frames within 5 pixels were joined to create tracks of the moving particles. To separate swimming bacteria tracks from other particles, tracks were only retained that had greater than 1-2 std above the mean deviation in position (Fig. 5A).

Though this automated software was useful for measuring swimming velocities of bacteria over short distances and could yield high n-values (sometimes in the range of 200-400 tracks per experiment for clear data with low background), we found it did not have high enough fidelity for use in chemotaxis assays, which requires long tracks, because the bacteria intersect and often dip in and out of the focal plane and thus a good intensity threshold value that separates the particle movement from noise cannot be obtained despite that the bacteria’s swimming is clearly discernible by eye. We therefore used the manual tracking addon within ImageJ for tracking swimming in chemotaxis assays (Fig. 5C-E, Fig. S3). This method retained high fidelity but was far more labor intensive, and so the number of bacteria tracked per experiment was between 30-60, equal to or higher to that performed for using similar assays from other studies (Behrens et al., 2016; Collins et al., 2016). We also include representative recordings from our experiments in which the decrease of swimming reversals and shift to smooth swimming is visible by eye (Fig. S3).

#### Quantification of H. pylori Swimming Reversals

Bacterial tracks as XY-coordinates from chemotaxis assays were input into an Excel spread sheet to quantify swimming reversals by calculating vector changes along the swimming trajectory. Here, we define a reversal as being a direction change of >300° with positions between frames at least deviating by 1.65 μm (to avoid reversals being counted when the bacteria is essentially motionless) (Fig. 5B). We found these to be reliable thresholds for replicating results based on visual analysis and removed a potentially large source of observer bias by applying the same standards to all samples analyzed.

#### Quantification of H. pylori Chemoattraction to Point Sources of Bleach

Data from micropipette chemotaxis assays were quantified using ImageJ’s Particle Analysis function (Fig. 6A-E). The number of bacteria in the field of view for each frame of data was counted and averaged over 2 s intervals (60 frames). These measurements were then normalized by dividing each count by the average number of bacteria observed over the five seconds immediately prior to insertion of the micropipette and treatment to obtain a plot of “Bacteria %” change over time (Fig. 6B,D-E). In these plots the average bacteria count pre-treatment is set to be 100 %, and each 2 s interval data point is divided by the average pre-treatment count. Videos of treatments are shown in Fig. S4. For assays with the *tlpABC* mutant that resulted in less substantial chemoattraction we performed additional analysis to help determine if the response was significant (Fig. S4C). We approximated chemoattraction to a newly established microgradient to be a linear process and fit bacteria counts over 1560 frames of post-treatment video to a simple linear model for NaOCl and buffer treatments. A significant positive correlation for NaOCl treatment was found (p = <0.00001) compared to buffer (p = 0.997), supporting that TlpD is sufficient for chemoattraction to bleach.

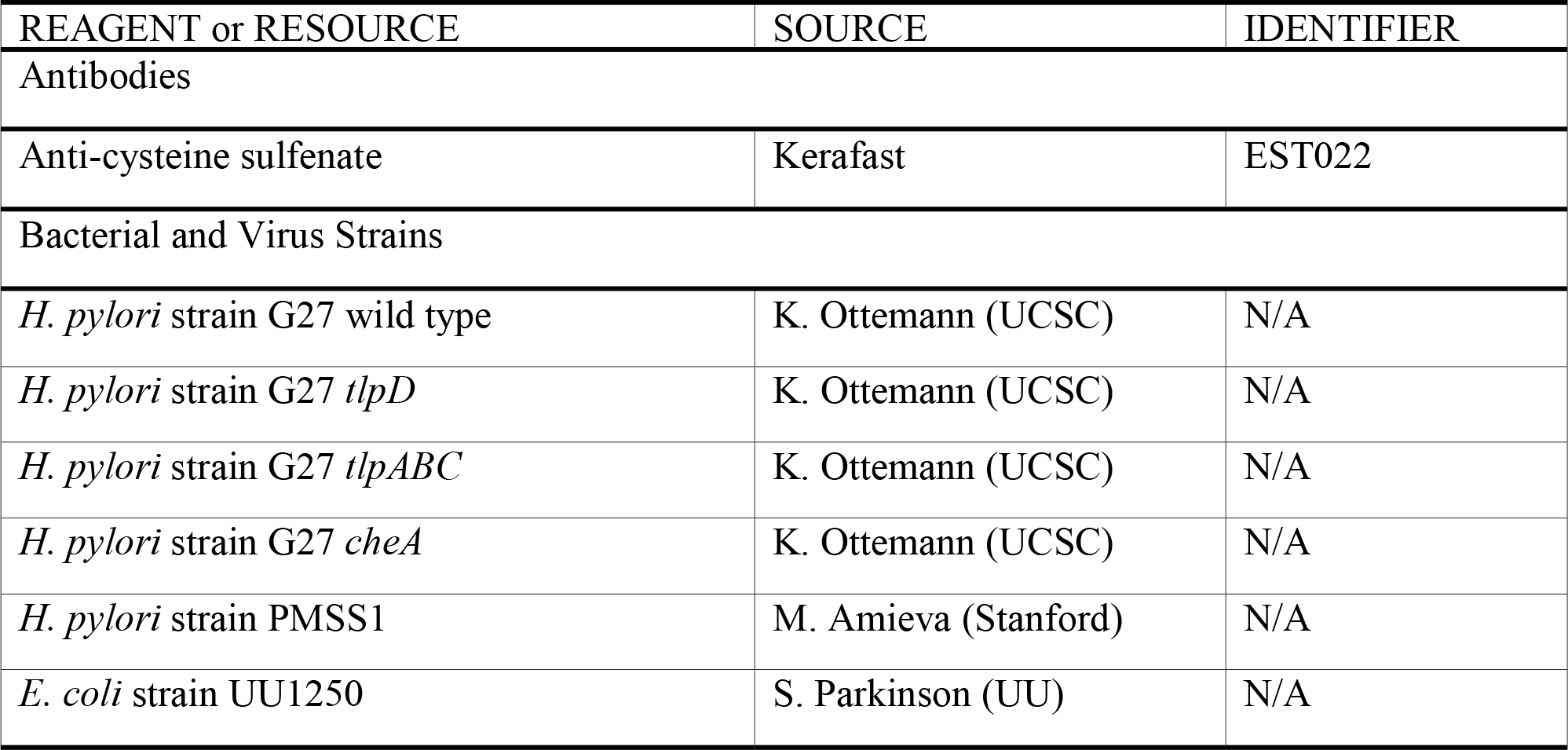
Key Resources Table.

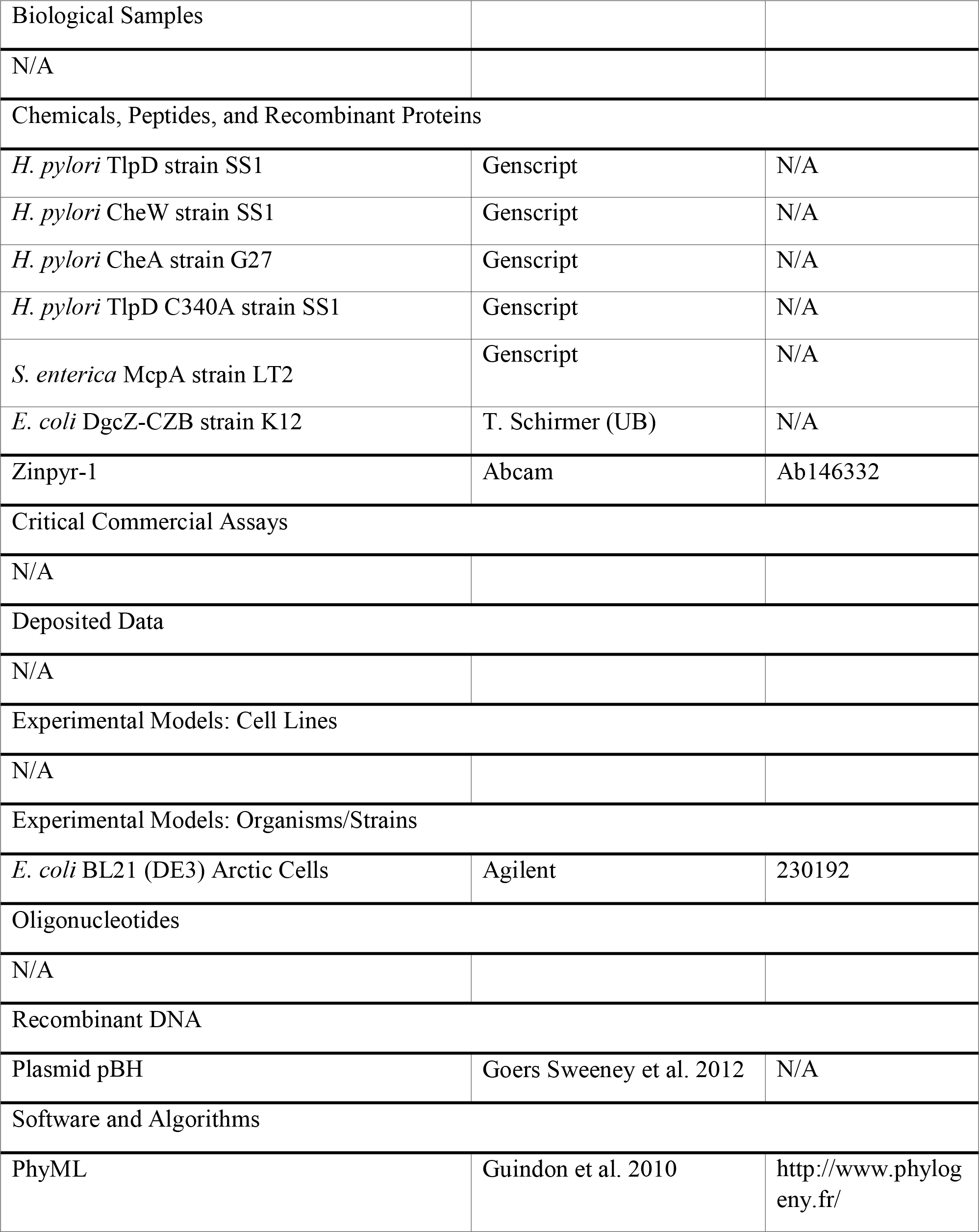

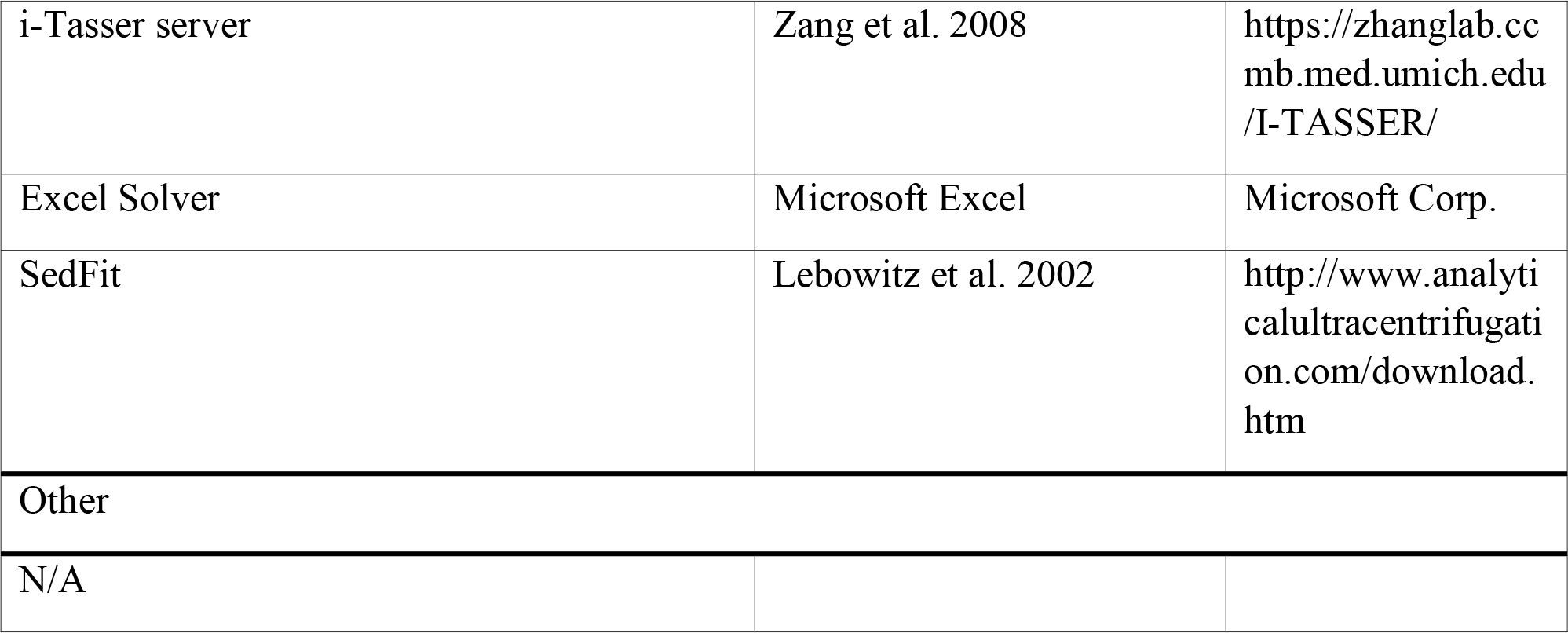

