## Supporting Information for "Helicobacter pylori Senses Bleach as a Chemoattractant Using a Cytosolic Chemoreceptor"

SUPPLEMENTAL INFORMATION

SUPPLEMENTAL FIGURES

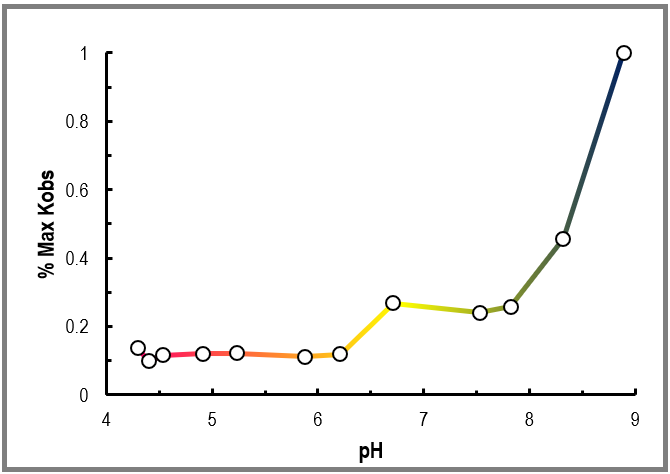

**Fig. S1 related to Fig 3.** Functional Assays Conducted at Various pH. Shown are results from a series of functional assays showing relative rates of CheA autophosphorylation for a titration from low (pink), neutral (yellow), and basic (blue) pHs. Experiments were run with 1 mM ATP, 4 µM CheA, 8 µM CheW, and 24 µM TlpD with 10 mM MgCl_2_, 100 mM NaCl and 200 mM of total buffer comprised of a combination of sodium citrate and tris. These data are opposite of that predicted to occur if TlpD was involved in direct sensing of pH in *in vivo* experiments, suggesting its role in acid sensing is indirect, and the higher activity at basic pH may be due to that CheA is more active at alkaline pH (see Method Details).

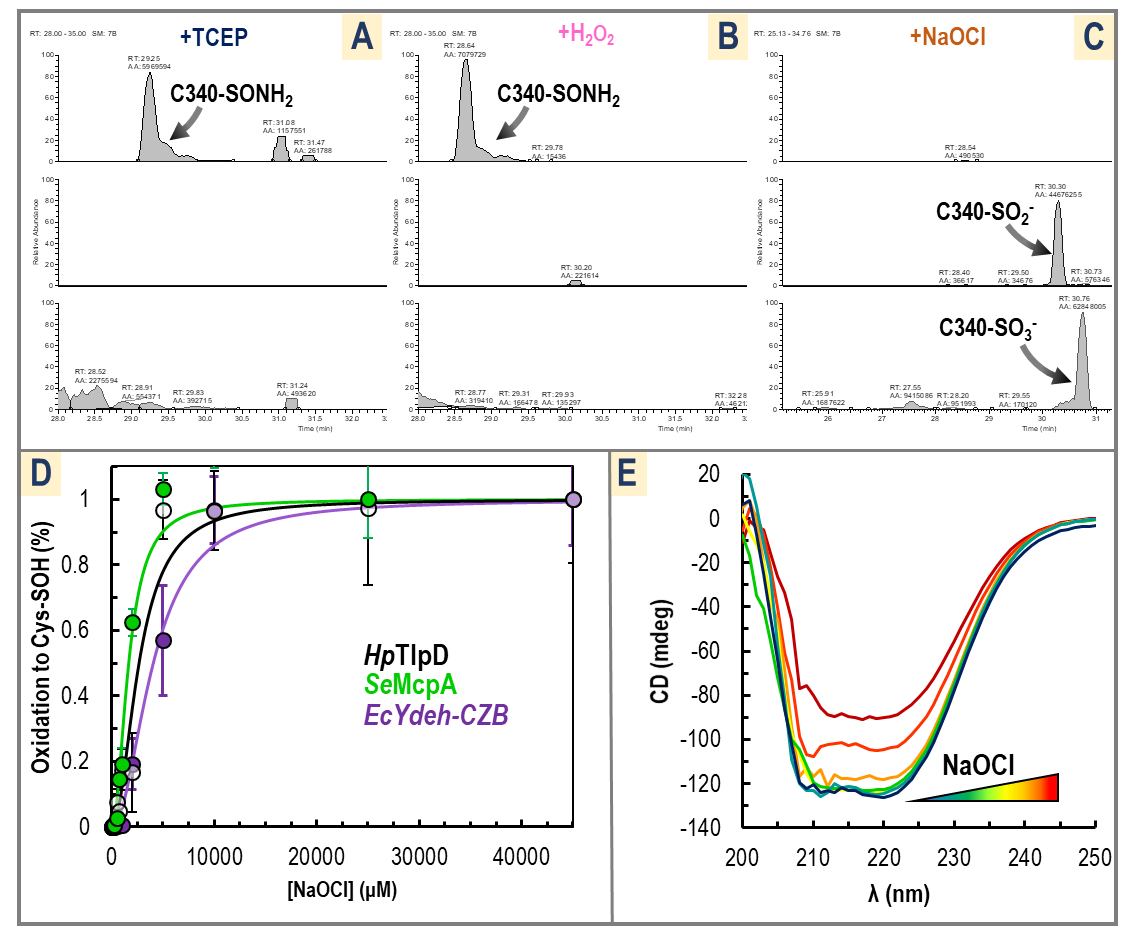

**Fig. S2 related to Fig 4.** Analysis of CZB Domain Oxidation by HOCl. (A-C) Analysis by MS/MS shows TlpD C340 is Oxidized by Bleach but not by Hydrogen Peroxide. Shown are MS/MS ion extractions for the TlpD C340-containing peptide NCRLGKWYYEGAGK from samples treated with 1 mM TCEP (A), hydrogen peroxide (B), or HOCl (C). For each sample, ion extractions for +3 charged peptides containing modifications of either alkylation are shown top (C340-SONH_2_, unreacted cysteine thiols modified by iodoacetamide), oxidation of C340 to a cysteine sulfinate (C340-SO_2_^-^) are shown middle, or oxidation of C340 to a cysteine sulfonate (C340-SO_3_^-^) are shown bottom on a fixed scale. Integrations values for peaks are noted with “AA.” The experiment and analysis are described further in the Material Details. (D) CZB protein domains from different bacterial species retain reactivity toward HOCl. Shown are reactions of purified *Hp*TlpD (black line, open circles), *Salmonella enterica* McpA (green), and the CZB domain of *E. coli* DgcZ (also called YdeH) with various concentrations of buffered HOCl. Solid lines are fits of the data to the Hill equation with a coefficient of 2, and markers shown are the average of triplicate independent measurements. Error bars are the sample standard deviation. (E) Shown is CD spectra of 20 µM TlpD treated for 1-H with buffer, 10 µM, 100 µM, 1 mM, 2 mM, 5 mM, and 10 mM NaOCl (dark blue, teal, green, yellow, orange, red, dark red, respectively). The protein remains folded even when treated with high concentrations of HOCl; signal deterioration for measurements with millimolar NaOCl corresponded to high electrical conductivity and is likely due to increased salt from the addition of NaOCl and not protein denaturation, as the curve shape remains similar (Greenfield, 2006). Sample buffer contained 25 mM NaCl, 20 mM Tris pH 7 and experiments were conducted at 20° C.

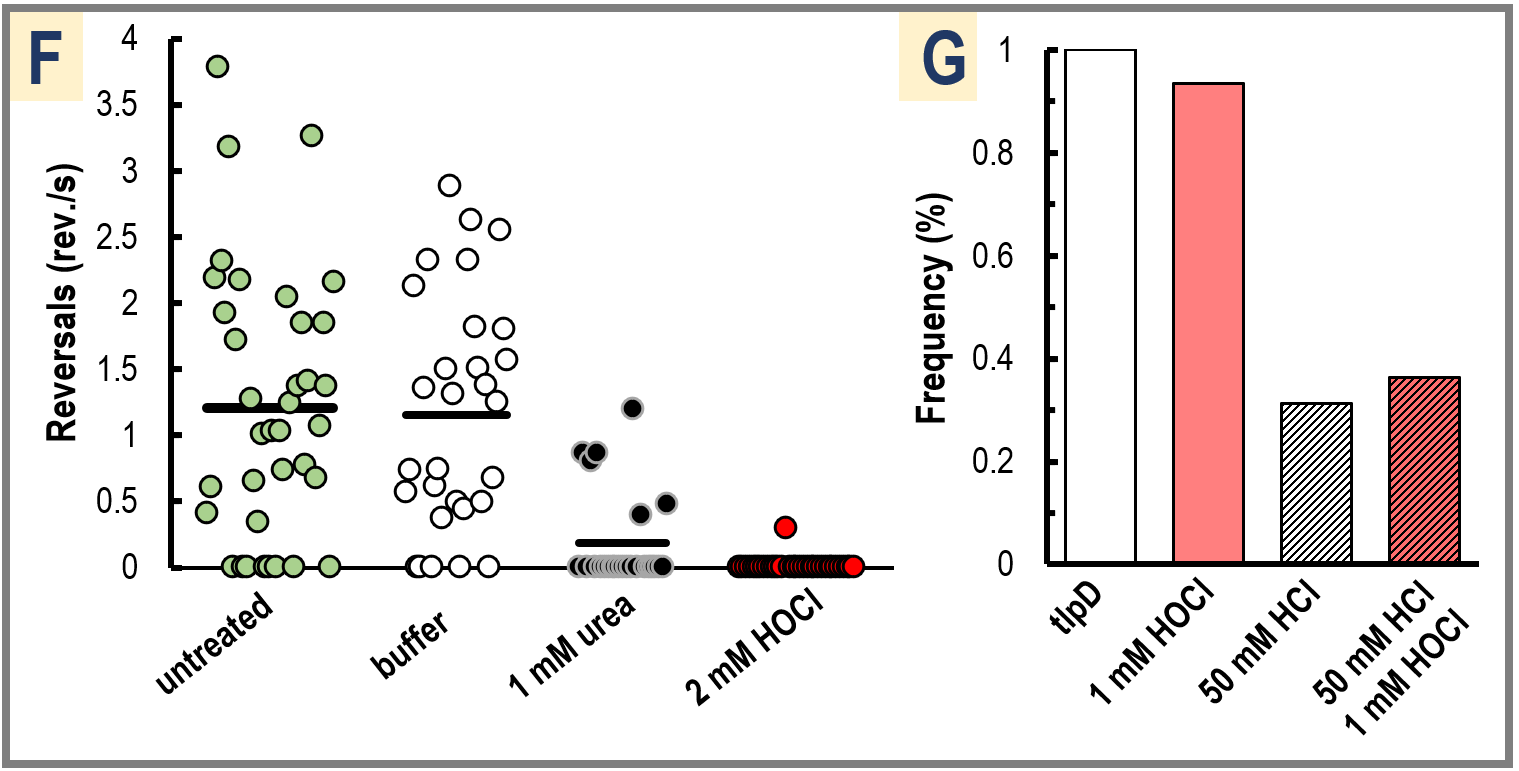

**Fig. S3 related to Fig. 5.** *H. pylori* shows a TlpD-dependent Decrease in Swimming Reversals in Response to HOCl. (A-C) Representative videos of wild type *H. pylori* strain G27 swimming shows behavior prior to treatment (A) <https://youtu.be/T6mr4gsFaWI>, treatment with a buffer solution of 50 mM Tris pH 7 in BB_10_ media (B) <https://youtu.be/eP-kvAlDWh4>, and following treatment with 4 mM HOCl in buffer (C) (see Materials & Methods) <https://youtu.be/_JinP69PpHg>. The video shown in A is the same swarm shown in C immediately before treatment. By default, the bacteria exhibit frequent swimming reversals, but treatment with bleach decreases reversal rates and leads to the chemoattraction behavior of smooth swimming. (D-E) Representative videos are shown of a triple chemoreceptor knockout *tlpABC* mutant (constructed in the G27 strain background) that possesses TlpD as its only chemoreceptor in response to buffer treatment (D) as in B <https://youtu.be/ubGE4OmFDL4>, and 4 mM HOCl treatment (E) as in C <https://youtu.be/Ses9lc4SgCc>. The *tlpABC* mutant shows a similar decrease in reversals and conversion to smooth swimming as seen for wild type. Recordings are shown at live speed. Note that treatments dilute the volume of cells by half, however we did not observe any significant change in reversal rates between untreated and buffer-treated samples at these low O.D.s (see panel F). For additional details on treatment protocol, reversal quantification, and microscopy see the Method Details. (F) Reversal rates for individual bacterial tracks representative treatments are shown, with untreated in gray-green, buffer-treated in white, 1 mM urea-treated in black, and 2 mM HOCl-treated in red. Black bars indicate the mean reversal rate. (G) Chemotaxis swarm assays with the *H. pylori* *tlpD* mutant (G27 background) are shown plotted as the frequency of the smooth swimming population. The *tlpD* mutant is almost entirely smooth swimming (white), and so further increases in smooth swimming by bleach addition cannot be observed in this way (light red). We therefore added 50 mM HCl, which can be sensed through TlpB and TlpA to increase reversals and decrease the smooth swimming population (white with black stripes). Cotreatment with 50 mM HCl and 1 mM HOCl (light red with black stripes) did not significantly decrease reversal compared to addition of HCl alone, consistent with bleach sensing being dependent on TlpD.

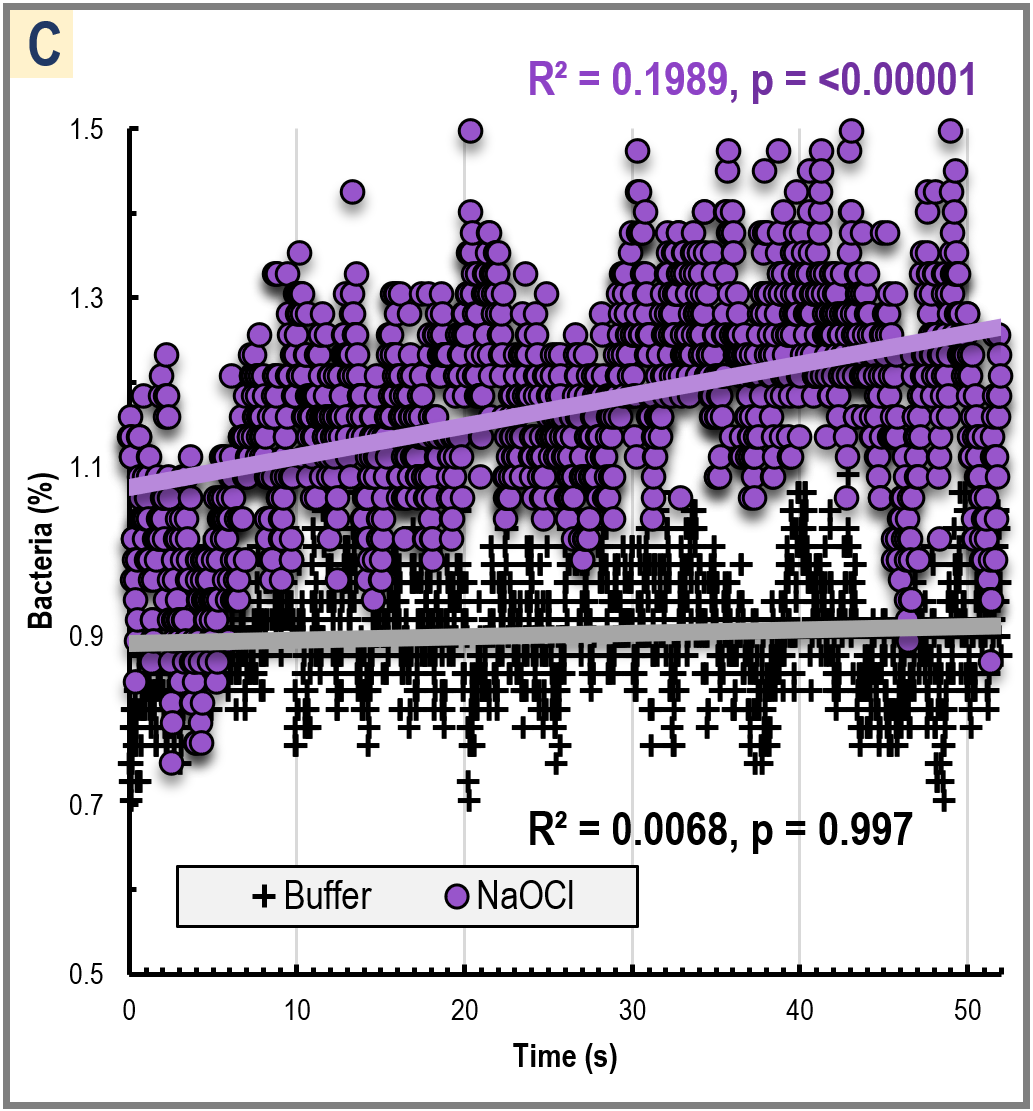

**Fig. S4 Related to Fig. 6.** TlpD is Required and Sufficient for *Helicobacter pylori* Chemoattraction to Bleach. (A-B) Representative videos from micropipette point-source assays are shown of wildtype PMSS1 <https://youtu.be/GW1Co3c5np8> and G27 <https://youtu.be/Y2Dt0Wk7eG8> *H. pylori* treated with either 500 mM sodium phosphate buffer pH 7 or 10 mM NaOCl diluted into buffer. Videos are shown in real time and begin 5 s prior to treatment, and then the micropipette is lowered into the swarm and treatment is applied over the next 30 s. (B) Videos are shown of *tlpD* <https://youtu.be/1s_ggO95ZYM> and *tlpABC* <https://youtu.be/NAEiBKCrZUE> mutants treated with buffer and NaOCl, as in A-B. (C) Statistical analysis of *tlpABC* point source experiments indicates a statistically significant positive correlation with NaOCl treatment (purple circles, p=0.1989, n=1560), and no statistically significant correlation for buffer treatment (black pluses, p=0.997, n=1560). Data shown are relative bacteria counts post-treatment for each frame of video (30 fps) normalized to 10 s prior to treatment. Linear models are fit to the post-treatment data and used for statistical analysis and comparison (see Method Details).

SUPPLEMENTARY TABLES

**Table S1.** Kinetic and Thermodynamic Properties of Reconstituted Signaling Complex.

| **Autophosphorylation^1^** | **Kinetics** |
| --- | --- |
| CheA | V_max ATP_ = 0.033 µM^-1^ min^-1^, K_M ATP_ = 136 µM |
| CheA, CheW | V_max ATP_ = 0.075 µM^-1^ min^-1^ |
| CheW, CheW, TlpD | V_max ATP_ = 0.43 µM^-1^ min^-1^ |
| **Complex Formation^2^** | **Dissociation Constants** |
| TlpD monomer ↔ dimer | K_D_ = 150 nM |
| CheA+CheW ↔ CheA-CheW | K_D_ = 14.6 µM |
| CheA-CheW+TlpD ↔ CheA-CheW-TlpD | K_D_= 15.2 µM |

1. Rates are for CheA autophosphorylation from data presented in Fig. 2 of either CheA alone, saturating concentrations of CheW, or maximally-activated by CheW and saturating concentrations of TlpD.
2. Parameters determined for complex formation are shown from data presented in Fig. 2. For titration of TlpD the concentration of CheA-CheW complex was estimated by titration of CheW against CheA (see Materials & Methods).
